# The microglia receptor protein TREM2 is essential for protective innate immune responses against herpesvirus infection in the brain

**DOI:** 10.1101/2023.03.16.532882

**Authors:** Stefanie Fruhwürth, Line S. Reinert, Carl Öberg, Marcelina Sakr, Marcus Henricsson, Henrik Zetterberg, Søren R. Paludan

**Affiliations:** Department of Rheumatology and Inflammatory Research, Institute of Medicine, Sahlgrenska Academy at the University of Gothenburg; Gothenburg, Sweden; Department of Psychiatry and Neurochemistry, Institute of Neuroscience and Physiology, Sahlgrenska Academy at the University of Gothenburg; Gothenburg, Sweden; Department of Biomedicine, Aarhus University; Aarhus, Denmark; Biomarker Discovery and Development, Research and Early Development, Cardiovascular, Renal, and Metabolism (CVRM), BioPharmaceuticals R&D, AstraZeneca; Gothenburg, Sweden; Clinical Neurochemistry Laboratory, Sahlgrenska University Hospital; Mölndal, Sweden; Department of Neurodegenerative Disease, UCL Institute of Neurology; Queen Square, London, United Kingdom; UK Dementia Research Institute at UCL; London, United Kingdom; Hong Kong Center for Neurodegenerative Diseases; Clear Water Bay, Hong Kong, China

## Abstract

Immunological control of viral infection in the brain is essential for immediate protection, but also for long-term maintenance of brain integrity. As the primary resident immune cell of the brain, microglia protect against viral infections through key macrophage functions, including release of the antiviral type I interferons (IFN-I) and clearance of infected cells. Microglia express the cytosolic DNA sensor cyclic GMP-AMP synthase (cGAS), which can bind viral DNA leading to signaling through stimulator of interferon genes (STING), and downstream immune activation. Here we report that herpes simplex virus (HSV) 1 infection of microglia leads to activation of IFN-I genes and pro-inflammatory cytokines. However, HSV1 also down-regulated expression of a subset of genes, including genes in the pathway engaged by the microglial receptor triggering receptor expressed on myeloid cells-2 (TREM2). Knockdown experiments revealed that TREM2 is important for viral activation of cGAS-STING signaling in microglia, induction of IFN-I, and phagocytosis of HSV1 infected neurons. Consequently, TREM2 depletion increased susceptibility to HSV1 infection in human microglia-neuron co-cultures and mice *in vivo*. Mechanistically, we show that TREM2 is essential for phosphorylation of STING, and downstream activation of the IFN-inducing transcription factor IRF3. We conclude that TREM2 is a novel component of the antiviral immune response in microglia, crucial for immediate host defense against HSV1 in the brain. Since both *TREM2* loss-of-function mutations and HSV1 serological status are linked to development of Alzheimeŕs disease (AD), this work opens the question whether defects in TREM2 could predispose to impaired viral clearance and post-infection pathological neurological changes.

## Introduction

Acute viral encephalitis is a severe disease, and often with fatal outcome if not treated (*1*). Herpes simplex encephalitis (HSE) is the leading cause of viral encephalitis in the Western world and is caused by herpes simplex virus type 1 (HSV1) infection (*2, 3*). HSV1 is a neurotropic human DNA virus that can reach the CNS via anterograde axon transport after infecting peripheral sensory neurons. Even though antiviral treatment is available for HSV1 and significantly improves survival rates, post-HSE is often associated with serious sequelae. Between 50 and 80% of the adult population are HSV1-seropositive (*4*), and although HSE develops only in very few cases, epidemiological data have linked HSV1 infection with development of Alzheimer’s disease (AD) (*5, 6*). Combined, the available data therefore suggests that immunological control of HSV1 infection in the CNS is important for immediate protection of the brain and for long-term maintenance of brain integrity.

The early innate immune response is crucial to restrict HSV1 spread in the CNS. Microglia are the primary resident immune cells of the brain and protect the brain parenchyma against pathogen invasions through key macrophage functions, such as the release of the antiviral type I interferons (IFN-I), cytokines, and phagocytosis of cell debris. Depletion of microglia increases viral spread in the CNS and consequently decreases the survival rates of mice infected with HSV1 (*7–9*). Optimal sensing and immune activation by microglia therefore require both continuous sampling of the local environment for danger signals, and pathogen sensing by a broad repertoire of innate pattern recognition receptors (PRRs).

For the sensing, microglia express high levels of many PRRs, including the cytosolic DNA sensor cyclic GMP-AMP synthase (cGAS). cGAS detects and binds to HSV1 DNA. leading to 2’3’cGAMP production and binding to stimulator of interferon genes (STING), which activates TANK-binding kinase 1 (TBK1) and interferon regulatory factor 3 (IRF3) (*10, 11*). Activated IRF3 translocates to the nucleus to activate the transcription of IFN-I genes, including IFNβ. Consequently, numerous IFN-stimulated genes (ISGs) are induced to exert direct antiviral activity through interference with specific steps in the viral replication cycle. Additionally, cGAS induces expression of inflammatory cytokines, such as IL6 and TNFα, via the NFκB pathway. We previously reported that cGAS and STING are highly expressed in microglia, which utilize this pathway to produce the bulk of IFN-I in the HSV1-infected mouse brain (*12*). In humans, functional deficits in TBK1 or IRF3 are associated with HSE susceptibility (*13, 14*), and several viral mechanisms to antagonize STING signaling have been identified (*15–18*). Collectively, this suggests that the cGAS-STING signaling axis plays an important role in control of HSV1 infection in the CNS.

For the sampling of the brain during both homeostatic and pathological conditions, microglia use a panel of receptors that bind a broad range of charged and hydrophobic molecules. These include for instance, scavenger receptor A-1, CD36, receptor for advanced glycation end products (RAGE), and triggering receptor expressed on myeloid cells-2 (TREM2). TREM2 is primarily expressed by microglia and interacts with a wide range of ligands including bacterial products, apoptotic cells, beta-amyloid (Aβ), anionic lipids, and apolipoprotein E (apoE) (*19*). Ligand engagement by TREM2 triggers downstream signaling through the adaptor protein DNAX activation protein 12 (DAP12) and recruitment and activation of the protein kinase SYK. In addition, TREM2 activity can be regulated via proteolytic processing of the receptor which releases a soluble TREM2 fragment (sTREM2) (*20*), the function of which remains to be fully elucidated. In humans, loss-of-function of TREM2 or DAP12 result in Nasu-Hakola disease, a serious disorder that is characterized by presenile dementia and bone cysts (*21, 22*). Partial loss-of-function of TREM2 increases the risk to develop AD several fold (*23, 24*).

Here we report that HSV1 infection of human microglia selectively down-regulates expression of genes in the TREM2 pathway. TREM2 was found to be important for viral activation of cGAS-STING signaling in microglia, and corresponding induction of the antiviral IFN-I response. Hence, TREM2 depletion in microglia and mice increased susceptibility to HSV1 infection in human microglia-neuron co-cultures and *in vivo*, respectively. These data identify TREM2 as a novel component of the antiviral immune response in microglia, which is actively targeted by HSV1.

## RESULTS

### HSV1 selectively blocks transcripts of the TREM2 pathway in hiPSC-derived microglia

We have previously reported that microglia play a central role in early sensing of HSV1 infection in the brain, and activation of host defense in mice (*12, 25*), and recently established that hiPSC-derived microglia are a powerful model system to study the antiviral immune response to herpesvirus infection (*18, 26*). The differentiation workflow from iPSCs to microglia and cortical neurons is illustrated in Figure S1A. Using bulk RNA sequencing (RNA-seq), we found that HSV1 infection profoundly altered microglial gene expression with a large proportion of the genes being affected (Figure S1B-C, Table S1). A broad panel of antiviral and proinflammatory genes were significantly upregulated (Fig. 1A-B). Gene ontology analysis suggests this response to impact on various processes, most notably antiviral response, leukocyte cell-cell interactions, and T cell activation (Fig. 1C). However, we also noted that a subset of the genes annotated to “response to virus” were downregulated (Fig. S1C), and this included a panel of ISGs, which were either down- or only modestly upregulated (Fig. S1D). To explore whether some of the downregulated genes were functionally related, we performed STRING analysis of the 200 most downregulated genes (Table S2). The analysis suggested four interaction nodes in this group of transcripts (Fig. 1D). One of these included several components of the TREM2 pathway, including *TREM2*, *TYROBP* (*DAP12*), *APOE*, and *SYK*. Collectively, HSV1 strongly activates antiviral transcription programs in human microglia, but also inhibits a subset of transcripts, including mRNAs encoding multiple components of the TREM2 pathway.

**Fig. 1.**
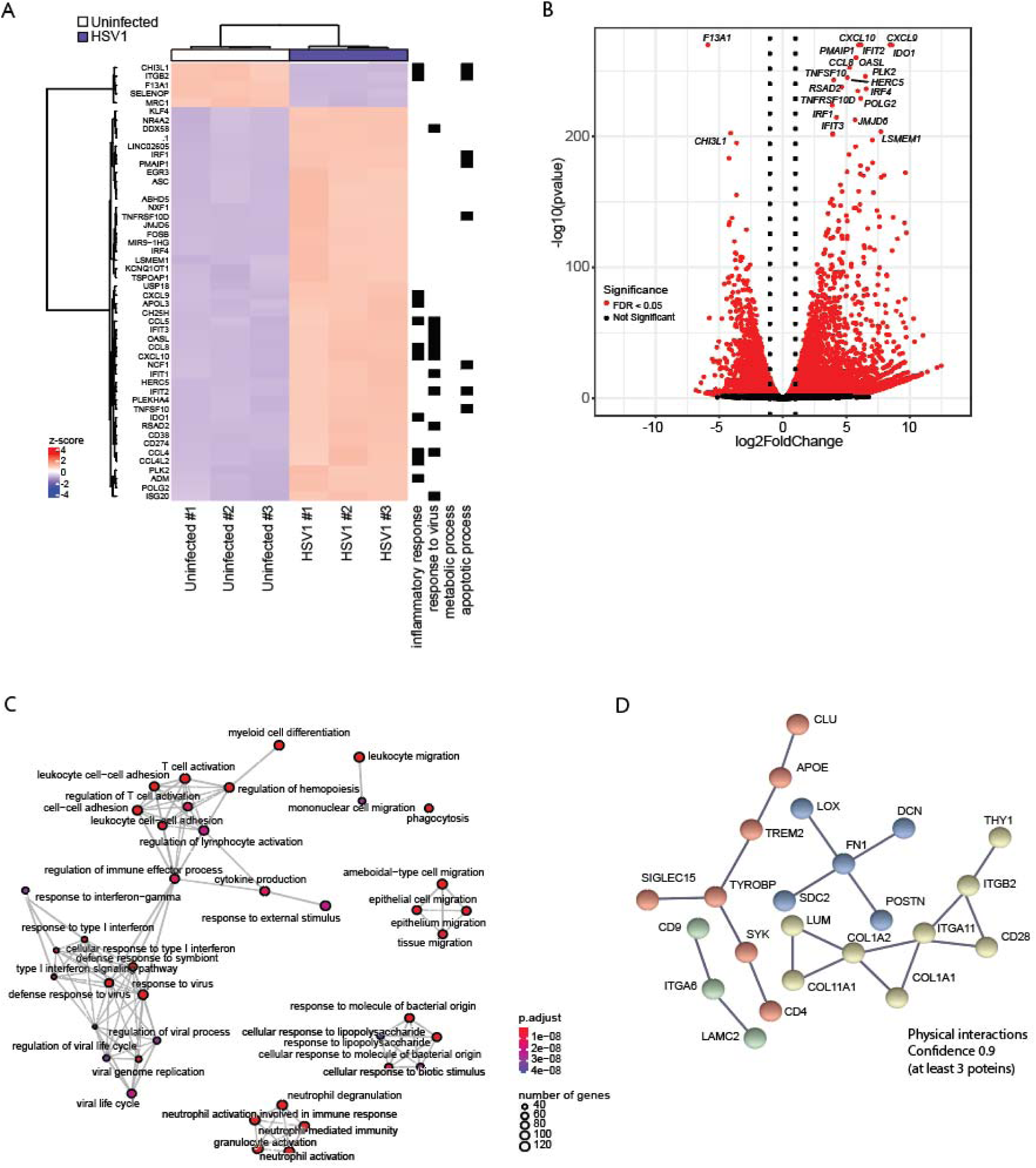
RNA-seq of HSV1-infected microglia. (**A-D**) HiPSC-derived microglia were infected with HSV1 (MOI 1) for 24 h. (**A**) Hierarchical clustering of genes differentially expressed in HSV1- versus uninfected microglia. Top 50 genes are shown. To the right is shown biological function annotated with the regulated genes. (**B**) Volcano plot of differentially expressed genes in uninfected versus HSV1-infected microglia (red, false-discovery rate (FDR) < 0.05). The top 20 differentially regulated genes are indicated. (**C**) Gene ontology enrichment plot in the category biological processes of the top 50 differentially expressed. (**D**) Network representation of the protein-protein interactions of the 200 most downregulated genes using STRING analysis. Nodes are represented with different colors for each node.

### HSV1 infection downregulates microglial TREM2 expression in hiPSC-derived microglia

We confirmed that TREM2 expression is significantly reduced in HSV1-infected microglia (Fig. 2). The levels of cellular full-length TREM2 protein (mature and immature form) as well as both of the proteolytic cleavage TREM2 products, *i.e.*, the C-terminal fragment (CTF) and secreted sTREM2, decreased substantially in HSV1 infected microglia (Fig. 2A-C). *TREM2* mRNA decreased in a virus dose-dependent and time-dependent manner (Fig. 2D-E). We observed this down-regulation using different HSV1 strains and in microglia differentiated from a separate iPSC line (Fig. S2A-C). To characterize the viral requirements for down-regulation of TREM2 expression, we treated virus with UV-light prior to infection or treated cells with acyclovir, which blocks viral DNA synthesis. Acyclovir did not alter HSV1-induced *TREM2* downregulation (Fig. 2F), whereas UV-inactivated HSV1 had lost the ability to alter *TREM2* expression levels (Fig. 2G). This suggests that TREM2 regulation requires viral activities occurring prior to viral DNA replication, which is thought to be limited in microglia, compared with more permissive cell types. The virion host shutoff gene (UL41) of HSV1 encodes a virion component that induces degradation of specific host mRNAs. We observed only a slight rescue of *TREM2* expression when using the ΔUL41 HSV1 mutant compared with wildtype HSV1 (Fig. S2H) suggesting that RNA degradation via this pathway is not the major mechanism exerting TREM2 downregulation.

**Fig. 2.**
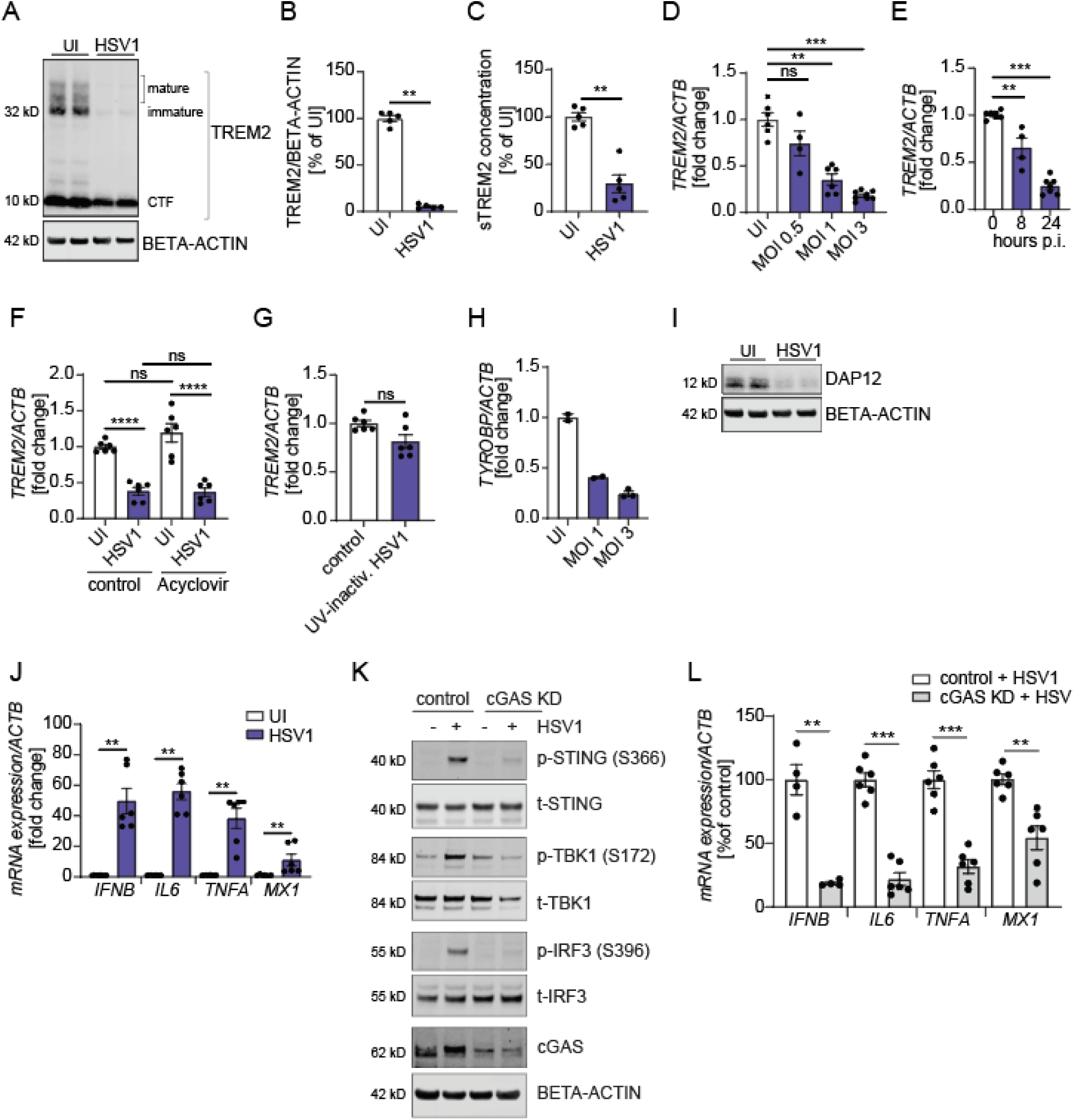
HSV1 downregulates microglial TREM2 expression. (**A**-**C**) HiPSC-derived microglia were infected with HSV1 (MOI 1) for 24 h. (**A**) A representative immunoblot of full-length TREM2 (TREM2) and C-terminal fragment (CTF) is shown. (**B**) Quantification of full-length TREM2 in uninfected (UI) microglia versus microglia infected with HSV1 (MOI 1) for 24 h. (**C**) sTREM2 levels were measured in cell supernatants of UI microglia versus microglia infected with HSV1 (MOI 1) for 24 h by ELISA. (**D**) *TREM2* mRNA levels were analyzed 24 h post-infection with HSV1 at different MOI’s as indicated. (**E**) *TREM2* mRNA levels were analyzed 8 and 24 h post-infection with HSV1 (MOI 1). (**F**) *TREM2* mRNA levels were analyzed 24 h post-infection with HSV1 MOI 1 with and without 50 µM Acyclovir. (**G**) HSV1 was UV-inactivated for 15 min. *TREM2* mRNA levels were analyzed 24 h after incubation with and without UV-inactivated HSV1 (MOI 1). (**H**) *TYROBP* mRNA levels were analyzed 24 h post-infection with HSV1 at different MOIs as indicated. (**I**) DAP12 protein levels were analyzed 24 h post-infection with HSV1 (MOI 1). A representative immunoblot is shown. (**J**) *IFNB*, *IL6*, *TNFA*, and *MX1* mRNA levels were analyzed in UI microglia versus microglia infected with HSV1 (MOI 1) for 24 h. (**K**-**L**) cGAS was knocked down in hiPSC-derived microglia using siRNA. Cells were used for experiments 4 days after transfection. (**K**) Control and cGAS KD microglia were infected with HSV1 (MOI 3) for 5 h. Representative immunoblots for the cGAS-STING signaling pathway and cGAS are shown. (**L**) *IFNB*, *IL6*, *TNFA*, and *MX1* mRNA levels were analyzed in control versus cGAS KD microglia 24 h post-infection with HSV1 (MOI 1). All figures represent at least 2-3 independent experiments; data are presented as mean ± SEM; P values were calculated by one-way ANOVA with Tukey’s multiple comparisons test (F) and Mann-Whitney test (B-E, G-H, J, L). ** P < 0.001; *** P < 0.0005; **** P < 0.0001.

In contrast to TREM2, and conforming the RNA-seq data, several other central microglial genes were not affected (*MERTK*, *AXL*) or upregulated (*TMEM119*, *P2YR12*) during HSV1 infection (Fig. S2D-E and S2F-G). Furthermore, we confirmed a significant downregulation of DAP12 on mRNA and protein level (Fig. 2H-I), thus validating the RNAseq data showing viral targeting of several factors in the TREM2 pathway. HSV1 infection strongly induced *IFNB*, *IL6*, *TNFA*, and the ISG *MX1* in hiPSC-derived microglia (Fig. 2J). We have previously shown in mice that microglia sense HSV1 infection in the CNS through the cGAS-STING pathway to activate the antiviral program including type I IFNs (*9, 12*). Here, we identified that this is also the case in human microglia. Knockdown of *cGAS* substantially impaired HSV1 sensing as shown by decreased phosphorylation of STING, TBK1, and IRF3 in response to HSV1 infection (Fig. 2K and S3D). Consequently, stimulation of *IFNB*, *IL6*, *TNFA*, and *MX1* was markedly impaired (Fig. 2L). Taken together, we observed robust downregulation of TREM2 protein and mRNA in HSV1-infected human microglia and show that the cGAS-STING pathway is essential for activating the antiviral program in human microglia. Based on these observations, we hypothesized that TREM2 might be actively involved in the innate antiviral immune response.

### Depletion of microglial TREM2 expression impairs the innate immune response to HSV1 infection in hiPSC-derived microglia

To determine the role of TREM2 during early HSV1 infection in hiPSC-derived microglia, we depleted *TREM2* using RNAi. The knockdown efficiency for full-length TREM2 was around 75 % (Fig. 3A-C and S3A) and we found sTREM2 to be significantly decreased as well (Fig. S3B). We did not observe a difference in cell viability in TREM2 KD cells compared with controls (Fig. 3D). To determine whether the innate immune response to HSV1 infection is affected by TREM2 knockdown (TREM2 KD), we measured mRNA levels of *IFNB*, *IL6*, *TNFA*, and *MX1* (Fig. 3E). Strikingly, we observed a significantly decreased innate immune response in TREM2-depleted microglia. Similar results were found with microglia differentiated from a separate iPSC line (Fig. S3C). Accordingly, we found impaired activation of the cGAS-STING pathway as shown by decreased phosphorylation of STING, TBK1, and IRF3 (Fig. 3F and S3D). Since sTREM2 has been suggested to exert biological functions such as protection against amyloid pathology (*27*), we next tested whether replenishing sTREM2 might be able to compensate for the loss of TREM2. Stimulation with sTREM2 had no effect on *IFNB* or *MX1* but induced the pro-inflammatory cytokines *IL6* and *TNFA* (Fig. S4A-C) as shown previously (*28*). Likewise, in the presence of HSV1 infection, we observed only a small increase in *TNFA* (Fig. S4D) suggesting that the decrease in sTREM2 during infection did not account for the effects observed.

**Fig. 3.**
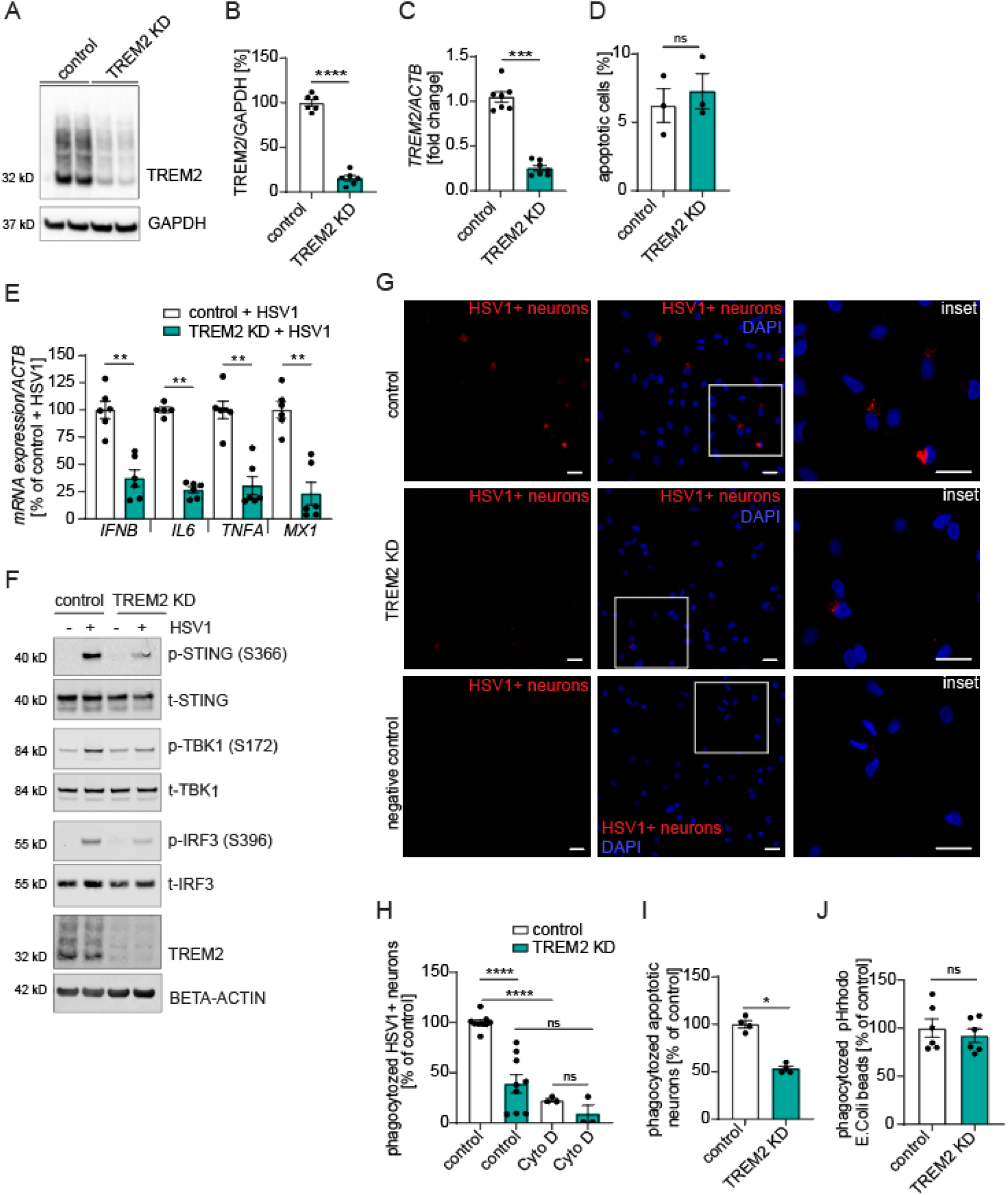
Depletion of TREM2 expression in microglia impairs innate immune responses to HSV1 *in vitro*. (**A**-**G**) TREM2 was knocked down in hiPSC-derived microglia using siRNA. Cells were used for experiments 4 days after transfection. (**A**) A representative immunoblot of full-length TREM2 (TREM2) of control and TREM2 KD microglia is shown. (**B**) Quantification of full-length TREM2 protein in control versus TREM2 KD microglia. (**C**) Quantification of *TREM2* mRNA in control and TREM2 KD microglia. (**D**) Apoptotic cells in control and TREM2 KD cultures were stained using an Annexin V antibody and quantified by flow cytometry. Quantification of apoptotic cells is shown. (**E**) *IFNB*, *IL6*, *TNFA*, and *MX1* mRNA levels were analyzed in control versus TREM2 KD microglia 24 h post-infection with HSV1 (MOI 1). (**F**) Control and TREM2 KD microglia were infected with HSV1 (MOI 3) for 5 h. Representative immunoblots for the cGAS-STING signaling pathway and TREM2 are shown. (**G, H**) HiPSC-derived neurons were infected with HSV1 (MOI 1) for 24 h, stained with a fluorescent cell tracker and added to control and TREM2 KD microglia for 15 h. (**G**) Cells were fixed and imaged using a confocal microscope (red=HSV1-infected neurons; blue=DAPI). The negative control were untreated cells. Scale bar = 20 µm. (**H**) Cells that had phagocytosed HSV1-infected neurons were quantified by flow cytometry (PE channel). Cytochalasin D (Cyto D, 5 µM) was added to specifically block phagocytosis. (**I**) Control and TREM2 KD microglia were incubated with fluorescent cell tracker-stained apoptotic neurons for 15 h. Cells that had phagocytosed apoptotic neurons were quantified by flow cytometry (PE channel). (**J**) Control and TREM2 KD microglia were incubated with pHrodo-*E. coli* beads for 3 h. Phagocytosis of pHrodo-*E. coli* beads was quantified using a microplate reader. All figures represent 2-3 independent experiments; data are presented as mean ± SEM; P values were calculated by Mann-Whitney test. * P < 0.05; ** P < 0.001; *** P < 0.0005; **** P< 0.0001.

Microglia protect against viral infection also via phagocytosis of infected cells, especially neurons. To determine whether phagocytosis was impaired in TREM2-depleted microglia, we analyzed microglial phagocytosis of HSV1-infected (HSV1+) neurons. Indeed, addition of fluorescently labeled HSV1-infected neurons to control microglia led to a prominent vesicular staining pattern (Fig. 3G, upper panel), whereas much less staining and fewer phagocytosing cells were observed in *TREM2* KD microglia (Fig. 3G, middle panel). No signal was observed in the negative control (Fig. 3G, lower panel). We further quantified the phagocytosis of HSV1-infected neurons using flow cytometry and found a significant decrease in the percentage of phagocytosing cells in TREM2 KD microglia (Fig. 3H). In the presence of the phagocytosis inhibitor cytochalasin D, the phagocytosis of HSV1-infected neurons was almost completely blocked in both groups, indicating that phagocytosis is necessary for entry of infected neuron debris into microglia. In addition, decreased phagocytosis of apoptotic neurons was observed in TREM2 KD microglia (Fig. 3I), whereas the phagocytosis of *E. coli* beads did not differ between control and TREM2 KD (Fig. 3J). This suggests a substrate-specific deficit in the phagocytic ability of TREM2-depleted microglia which has been reported previously for TREM2 loss of function (*29*). These results show that the antiviral response in *TREM2*-depleted microglia is affected in several ways, including impaired IFNB induction via the cGAS-STING pathway, and clearance of infected neurons.

### TREM2 is essential for control of HSV1 infection *in vitro* and *in vivo*

HSV1 is a neurotropic virus, and replication predominantly occurs in neurons in the CNS. To investigate whether the impaired immune response in *TREM2* KD microglia affects the control of HSV1 infection *in vitro* we co-cultured hiPSC-derived cortical neurons and microglia (Fig. S1A). As reported previously in mice (*12*), we observed that human microglia are potent producers of type I IFN (Fig. S5A). Like in microglia cultures, *TREM2* mRNA as well as secreted sTREM2 protein levels were significantly decreased in HSV1-infected co-cultures (Fig. 4A and 4B). Interestingly, less viral replication and higher cell viability was observed in co-cultures compared with neurons alone (Fig. S5B and S5C). Importantly, significantly higher viral replication occurred in TREM2-depleted co-cultures, which was accompanied by decreased cell viability (Fig. 4C and 4D).

**Fig. 4.**
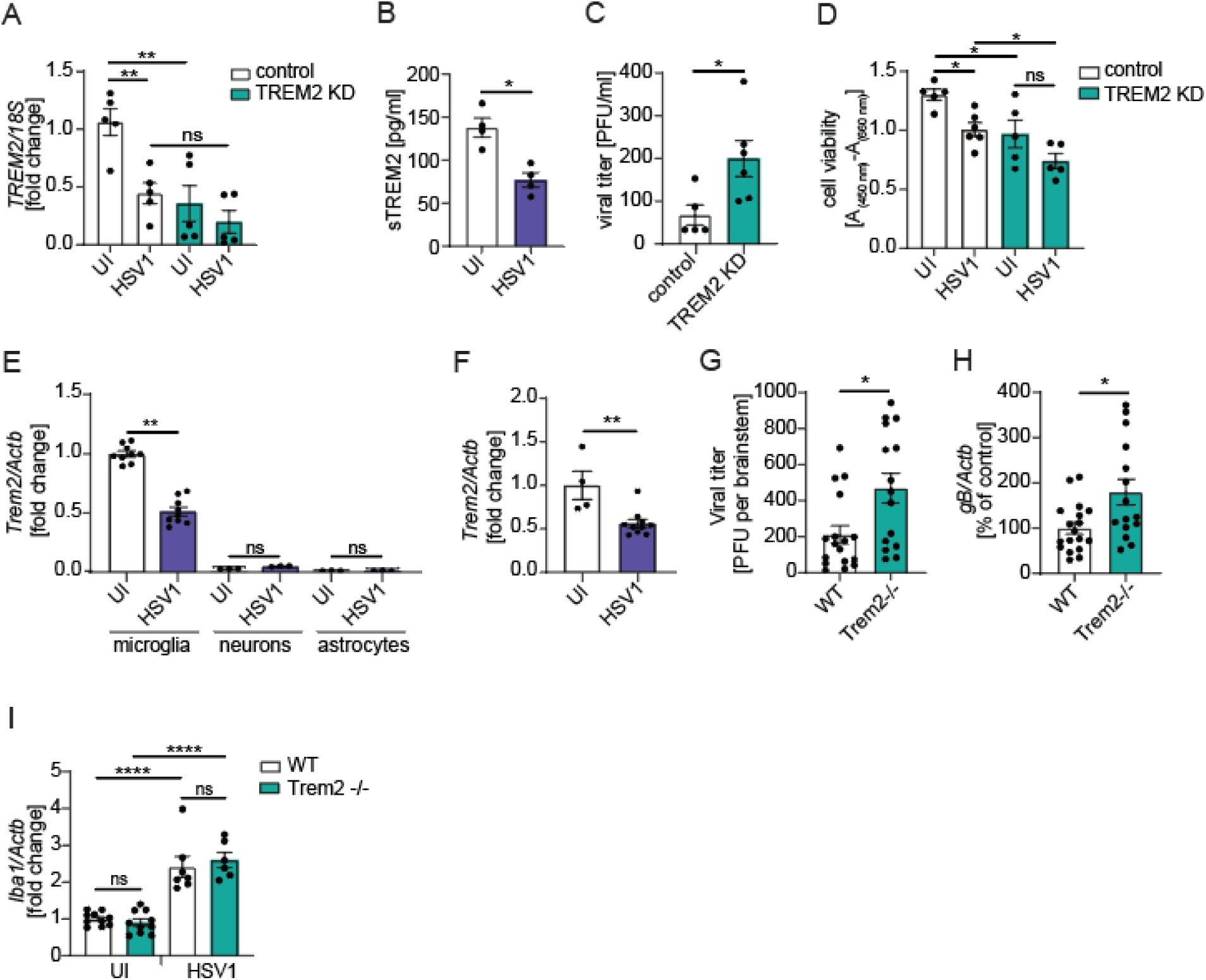
TREM2 is essential for control of HSV1 infection *in vitro* and *in vivo*. (**A**) *TREM2* mRNA levels were analyzed 24 h post-infection of hiPSC-derived co-cultures of microglia and neurons with HSV1 (MOI 1). (**B**) sTREM2 levels were measured in supernatants of co-cultures 24 h post-infection with HSV1 (MOI 1) by ELISA. (**C**) Viral titers in co-culture supernatants 24 h post-infection with HSV1 (MOI 1) were quantified by viral plaque assay. (**D**) Cell viability was measured in co-cultures 24 h post-infection with HSV1 (MOI 1) using the CyQUANT XTT assay. Specific absorbance, which was calculated by using the formula: A_(450 nm)_-A_(660 nm)_, is shown and reflects cell viability. (**E**) *Trem2* mRNA levels were analyzed in primary murine microglia, neurons, and astrocytes 6 h post-infection with HSV1 (McKrea, MOI 3). (**F**) *Trem2* mRNA levels were analyzed in WT mouse brainstem 4 days post-infection with HSV1 via the corneal route (2 × 10^6^ PFU/cornea). (**G**) Viral titers of isolated brainstems 4 days post-infection with HSV1 via the corneal route (2 × 10^6^ PFU/cornea) were quantified by plaque assay. (**H**, **I**) *gB* and *Iba1* mRNA levels were quantified in brainstem of WT and Trem2 -/- mice 4 days post-infection with HSV1 via the corneal route (2 × 10^6^ PFU/cornea). All figures represent 2-3 independent experiments; *n* = 15-18 per group of mice; data are presented as mean ± SEM; P values were calculated by one-way ANOVA with Tukey’s multiple comparisons test (A, D, I) and Mann-Whitney test (B, C, E-H). * P < 0.05; ** P < 0.001; **** P < 0.0001.

To explore whether the phenomenon observed in human microglia was also observed in mice, we infected mouse brain cells *in vitro* or live animals through the corneal route, which leads to infection in the brain stem. Importantly, we observed HSV1-induced *Trem2* downregulation in primary mouse microglia and in the brainstem of HSV1-infected wildtype (WT) mice (Fig. 4E and 4F). *Trem2* expression was not detected in astrocytes or neurons (Fig. 4E). To determine whether TREM2 is essential for control of HSV1 infection *in vivo*, we compared viral load in WT versus *Trem2^-/-^* mice. Consistent with the data from the human co-culture system, we observed significantly increased levels of infectious virus and viral mRNA in the brainstem of HSV1-infected *Trem2^-/-^* mice compared with WT (Fig. 4G and 4H). We did not detect differences in *Iba1* expression in the brain stem between genotypes suggesting that the observed infection phenotype was unlikely to be due to a difference in microglial content (Fig. 4I). Collectively, these findings suggest that TREM2 depletion leads to impaired control of HSV1 infection *in vitro* and *in vivo*.

### TREM2-depleted microglia respond normally to cGAMP

Since we observed impaired activation of the cGAS-STING pathway by HSV1 in TREM2-depleted microglia (Fig. 3F and S3D), we wanted to test the functionality of this viral sensing pathway. To this end, we stimulated microglia with cGAMP which directly activates STING. Interestingly, we found no significant difference in mRNA levels of *IFNB*, *IL6*, *TNFA*, and *MX1* in cGAMP-stimulated control versus TREM2-depleted microglia (Fig. 5A). Accordingly, we did not observe a difference in phosphorylation of STING, TBK1, and IRF3 in cGAMP-stimulated control versus TREM2-depleted microglia (Fig. 5B and S6A). Also, we found no difference in *cGAS* mRNA or cGAS protein expression between control and TREM2 KD microglia (Fig. 5C-D). These data suggest that TREM2 does not affect STING signaling per se. In addition, we observed that HSV1-induced cGAMP accumulation in microglia was not affected by knock-down of TREM2 (Fig. 5E), whereas STING phosphorylation was reduced (Fig. 3F). In uninfected cells, cGAMP was below detection limit (data not shown). Our results indicate that, while STING signaling alone is not dependent on TREM2, this microglial receptor augments HSV1-induced cGAS-STING signaling down-stream of cGAMP production, and upstream of STING phosphorylation.

**Fig. 5.**
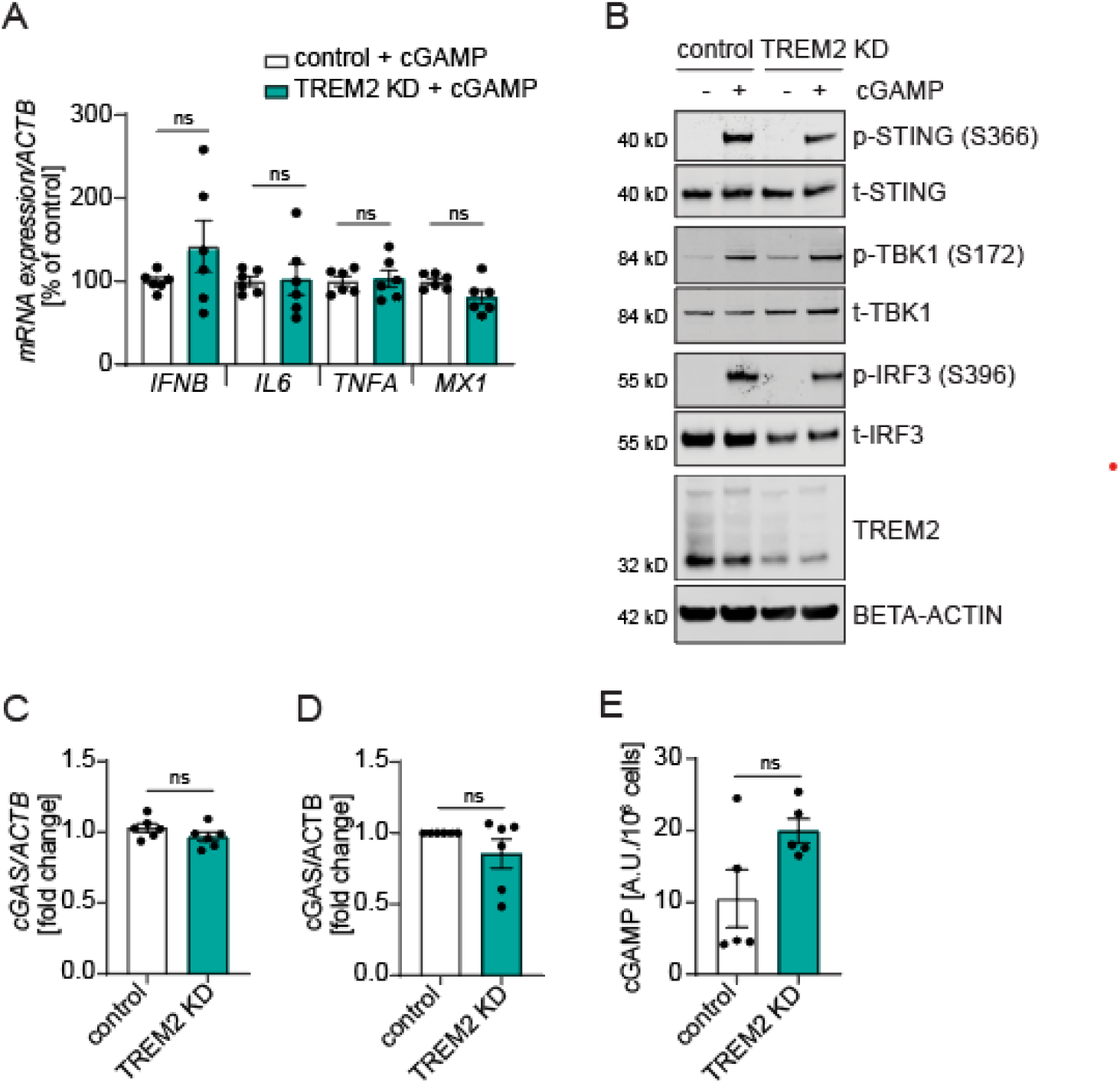
The antiviral program is not impaired in response to cGAMP in TREM2 depleted hiPSC-derived microglia. (**A-B**) Control and TREM2 KD microglia were stimulated with 100 µM cGAMP for 5 h. (**A**) *IFNB*, *IL6*, *TNFA*, and *MX1* mRNA levels are shown. (**B**) Representative immunoblots of the cGAS-STING axis and TREM2 are shown. (**C**) *cGAS* mRNA levels are shown in control and TREM2 KD microglia. (**D**) Quantification of cGAS protein levels in control and TREM2 KD microglia is shown. (**E**) Quantification of cGAMP levels in HSV1-infected control versus TREM2 KD microglia 24 h post-infection. *­­A.U.* arbitrary units. All figures represent 2-3 independent experiments; data are presented as mean ± SEM; P values were calculated by Mann-Whitney test.

### Modulation of TREM2 activity via SYK modulates the innate immune response to HSV1 infection in microglia

The protein kinase SYK, which is an immediate downstream target of TREM2, has been shown to be crucial for STING-mediated IFN induction by HSV1 (*30*). To determine whether TREM2 might modulate the antiviral immune response via SYK, we modulated TREM2 signaling and SYK activity. First, we used a TREM2-activating antibody (AF1828), which induces downstream signaling via SYK activation. Since microglia express Fc receptors, IgG was used as an isotype control. As previously reported (*31*), stimulation of human microglia with the AF1828 antibody rapidly induces SYK phosphorylation in control but not in TREM2 KD cells (Fig. 6A). Using this approach, we observed that stimulation with AF1828 amplified the early immune response to HSV1 infection as shown by significantly increased levels of *IFNB*, *IL6*, *TNFA*, and *MX1* (Fig. 6B). Next, we blocked TREM2 downstream signaling using a selective SYK inhibitor (ER27319) (Fig. 6C-D). In the presence of ER27319, SYK phosphorylation was substantially decreased upon stimulation with AF1828 (Fig. 6C). Consequently, we found significantly decreased levels of *IFNB*, *IL6*, *TNFA*, and *MX1* upon SYK inhibition in HSV1-infected microglia (Fig. 6D). In addition, we observed that HSV1 treatment itself induced a transient SYK phosphorylation in microglia (Fig. 6E-F). Collectively, these results show that SYK activation downstream of TREM2 augments the antiviral response in microglia.

**Fig. 6.**
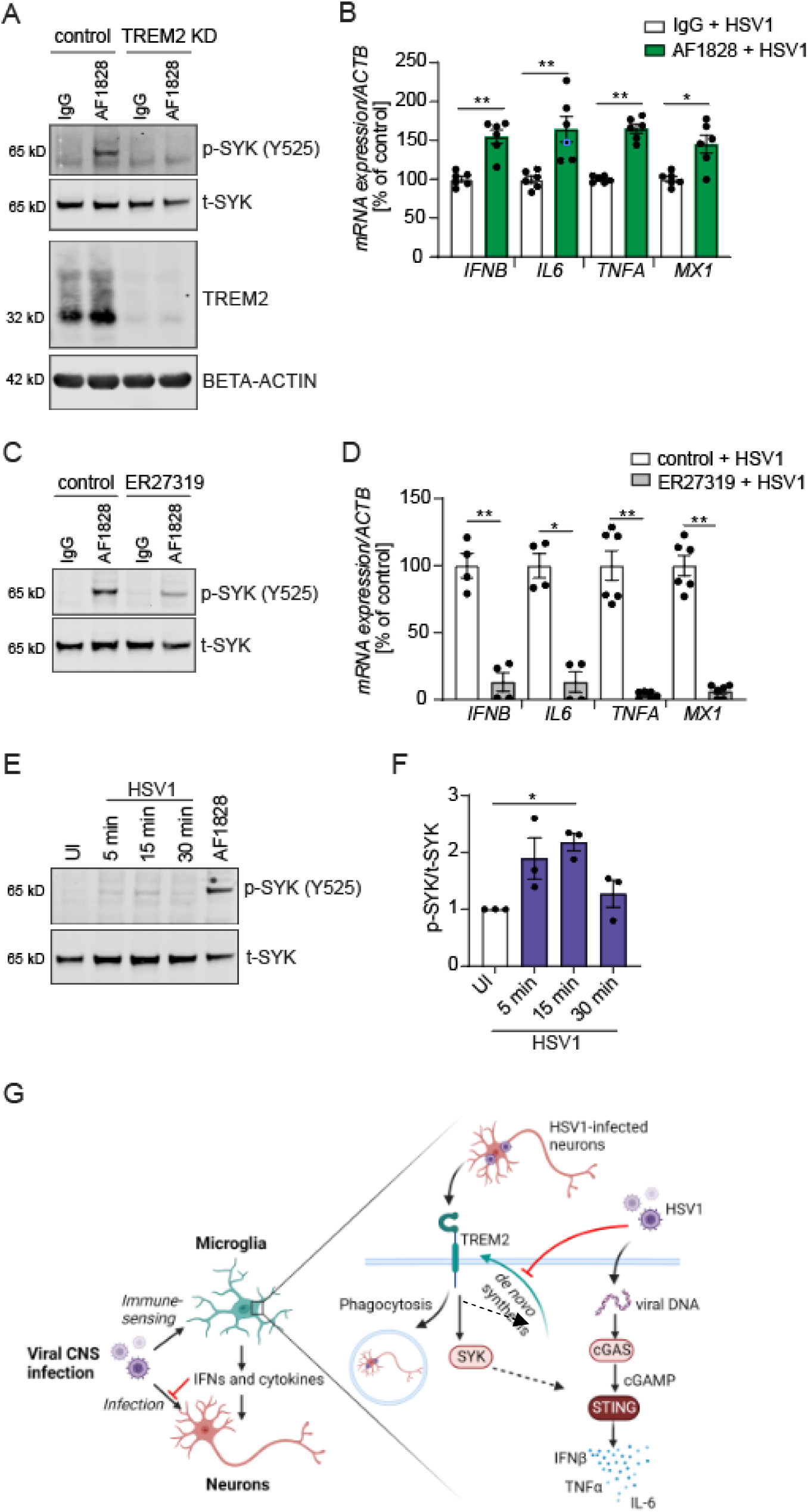
Modulation of TREM2 activity via SYK modulates the innate immune response to HSV1 infection in hiPSC-derived microglia. (**A**) Control and TREM2 KD microglia were incubated with 0.8 µg/ml control IgG or the TREM2-activating antibody AF1828 for 15 min. Representative immunoblots of p-SYK and TREM2 are shown. (**B**) *IFNB*, *IL6*, *TNFA*, and *MX1* mRNA levels were analyzed in microglia after incubation with HSV1 (MOI 1) in combination with control IgG or AF1828 for 8 h. (**C**) Microglia were incubated with 0.8 µg/ml control IgG or AF1828 and with or without 5 µM ER27319 (SYK inhibitor) for 10 min. A representative immunoblot of p-SYK is shown. (**D**) Microglia were infected with HSV1 (MOI 1) with and without 5 µM ER27319 for 8 h. *IFNB*, *IL6*, *TNFA*, and *MX1* mRNA levels are shown. (**E, F**) Microglia were infected with HSV1 (MOI 5) for the indicated time points or stimulated with 0.8 µg/ml AF1828 for 30 min. (**E**) A representative immunoblot is shown. (**F**) Quantification of p-SYK/t-SYK from three independent experiments is shown. (**G**) Illustration of the roles of microglial TREM2 during viral CNS infection. All figures represent 2-3 independent experiments; data are presented as mean ± SEM; P values were calculated by Mann-Whitney test (B, D) and one-way ANOVA with Tukey’s multiple comparisons test (F). * P < 0.05; ** P < 0.001.

## DISCUSSION

In this study, we identify TREM2 as an essential component of the microglial immune response to HSV1 and demonstrate that HSV1 actively downregulates this pathway through a mechanism activated early after viral entry into the microglia. Depletion of TREM2 expression impaired the innate immune response to HSV1 infection, including type I IFN induction, and consequently promoted viral replication in human brain cell cultures and in mice *in vivo*. These data further support the idea of the essential role for microglia in early antiviral defense and provide mechanistic insight to the virus-cell interactions that govern the outcome of HSV1 infection in the brain.

Using bulk RNA-seq, we found that HSV1 infection profoundly induced antiviral and proinflammatory genes but also inhibited a subset of transcripts, including multiple components of the TREM2 pathway. In support of a role for TREM2 in the antiviral response, we found a dramatic decrease in the HSV1-induced activation of Type I IFNs and ISGs in TREM2 KD microglia. Even though we found that HSV1 infection robustly downregulates microglial TREM2 expression within 24 hours, we observed significant differences between control and TREM2 KD microglia in terms of antiviral defense mechanisms. These results indicate that TREM2 is essential for the immediate response to infection, and that the virus seeks to target this response. This is in line with previous reports suggesting that the very early microglial immune response is crucial for control of HSV1 infection (*7–9, 12*). We did not explore other pathways, which were identified through the analysis of the RNAseq data to be altered. However, it is interesting that two integrin-involving pathways were modulated by the virus, and integrins are important for the ability of microglia to migrate to sites of damage in the brain (*32*).

As the primary immune cells of the brain, microglia recognize and remove injured and apoptotic cells. In our *in vitro* assay, phagocytosis of HSV1-infected neurons was found to be dependent on TREM2. This is similar to what has previously been reported for microglial phagocytosis of apoptotic neurons (*33–35*). HSV-infected neurons do not abundantly undergo apoptosis (*36*), and these data therefore illustrate how a range of different ligands can trigger TREM2-driven phagocytosis. In addition, the data from co-culture and *in vivo* studies identified TREM2 as an important player in the control of HSV1 infection and spread in the brain. The observed effects of TREM2 loss-of-expression are likely a combination of impaired innate immune response, decreased clearance of infected neurons as well as impaired recruitment of microglia to infected areas. It has been suggested that rapid recruitment of microglia is critical to limit viral spread (*8*) and TREM2 deficiency has been demonstrated to impair chemotaxis and microglial responses to neuronal injury (*37*). Whether this also applies to HSV1-infected neurons needs further investigation.

SYK is highly expressed by haemopoietic cells, and the activation of this kinase has recently been shown to be required for STING-mediated IFN induction by HSV1 in different cell lines (*30*), and also to drive STAT1 activation in response to infection with RNA viruses (*38*). In our experimental system with iPSC-derived human microglia we observed that HSV1 infection led to rapid phosphorylation of SYK, and treatment with the SYK inhibitor ER27319 diminished the antiviral IFN immune response to HSV1 infection, which we found mainly to proceed through the cGAS-STING pathway. We propose that TREM2 engagement on microglia by HSV1 or HSV1-infected cells triggers SYK activation, thus amplifying signaling through the cGAS-STING pathway by augmenting STING activity dependent on its phosphorylation on S366 (Fig. 6G). Upon viral entry into microglia this response was blunted by down-regulation of *TREM2* mRNA levels, which consequently impairs the immediate antiviral host response. We did not fully identify the mechanism through which HSV1 down-modulates levels of TREM2 transcripts, although we did find that it was dependent on viral gene expression, but independent of viral DNA replication. HSV1 can enter into many cell types, but only replicate productively to high titers in a narrower subset of cells. Our data demonstrate that it is not a “dead end” for HSV1 to interact with microglia, although these cells cannot be productively infected. Rather this interaction can impair the antiviral IFN-I response, and hence promote the replication of virus that has entered into more permissive cells, such as neurons.

HSV1 infection has been epidemiologically associated with development of AD, primarily in carriers of the *APOE ε4* allele (*39*). AD is the most common form of dementia and affects more than 10% of people over age 65 (*40*). The majority of genetic AD risk factors are implicated in microglial and innate immune cell functions (*41*), including a hypo-functional TREM2 allele which increases the AD risk several fold (*23, 24*). It is, therefore, tempting to speculate that the HSV1-induced downregulation of TREM2 might aggravate AD-related pathologies, especially amyloid plaque accumulation. Recent data highlight a role for TREM2 in activating and directing microglial responses in disease (*42*) and controlling the ability of microglia to surround, contain, or clear Aβ (*43, 44*). Thus, microglial activation plays beneficial roles in prevention of AD, and impaired microglial activation results in a significantly increased risk for AD. Given the growing interest in development of TREM2 activating antibodies as a potential therapeutic approach for AD (*45, 46*), novel insights into TREM2 function add valuable information. Immune activation to infection is essential for eventual control of virus infection in the brain. However, the brain damage that occurs during HSE is caused by both viral replication and excessive inflammatory response (*47*). Our results suggest that elevating TREM2 function might be beneficial during acute HSV1 infection in the brain, but also provide a possible mechanism for how viral reduction of TREM2 expression could affect Aβ pathology and accelerate disease processes.

The present work is mainly based on work in iPSC-derived microglia and neurons. Therefore, although this provides immediate demonstration that the observed phenomena occur in primary human cells, it does not show the relevance under natural infection in the human brain. This will require further studies, including exploration of the association of genetic variants of TREM2 with HSV1 infection in the brain. At the mechanistic side, it a limitation of our work that the molecular structure of HSV1 or HSV1-infected cells that trigger TREM2 in microglia, including SYK activation, augmentation of gene expression, and phagocytosis was not identified. We find that this is without the scope of this work and requires independent in-depth studies. Interestingly, it has been reported that for instance DNA, different lipids, and complex carbohydrates, which are present on viral particles and stressed cells, have been reported as TREM2 agonists (*19*).

Here, we demonstrate that the TREM2 pathway is downregulated by HSV1 infection, and that this innate immune pathway plays a central part of the early antiviral immune response to HSV1 infection in the brain. Therefore, specific immunomodulatory strategies combined with antiviral therapy represent a promising path for better control of this devastating disease and maintenance of brain integrity.

## MATERIALS AND METHODS

### Viruses and reagents

A neurovirulent clinical HSV strain was used for most *in vitro* experiments unless indicated differently: HSV1 2762 was isolated from the brain of a patient with fatal HSE during a clinical trial of acyclovir treatment (*48*). In addition, a KOS wildtype and a KOS mutant strain (ΔUL41) were used. For *in vivo* and primary murine cell *in vitro* experiments, the McKrae strain was used. The titers of the stocks used were determined by plaque assay on Vero cells (ATCC). AF1828 was from R&D, control IgG was from Santa Cruz (sc-2028), 2′-3′cGAMP was from InvivoGen, human recombinant sTREM2 was from Sino Biological, and Acyclovir was from Pfizer.

### Generation of human hiPSC-derived microglia

Two human iPSC lines WTSIi015-A (EBiSC through Sigma-Aldrich) and ChiPSC22 (Takara Bio Europe) were used. WTSIi015-A iPS cells were used for most experiments unless indicated differently. IPS cells were maintained on Matrigel (Corning) in mTeSR1+ medium (Stemcell Technologies). IPSC colonies were dissociated into single cells using TrypLE Express (Thermo Fisher Scientific). 4*10^6^ iPSCs were seeded per Aggrewell 800 (Stemcell Technologies) in a 24-well plate in 2 ml embryonic body medium (EBM). EBM consisted of mTeSR1+ medium supplemented with 10 μM ROCK inhibitor, 50 ng/mL BMP-4, 20 ng/mL SCF, and 50 ng/mL VEGF-121 (all from Peprotech). Cells were cultured for 4 days in Aggrewells to form embryonic bodies (EBs) with half media change (1 ml) every day. EBs were harvested using an inverted cell strainer (40 μm), and around 15 EBs were plated per 6-well in hematopoietic medium (HM). HM consisted of X-VIVO 15 medium (Lonza) supplemented with 2 mM Glutamax, 100 U/mL penicillin, 100 μg/mL streptomycin, 55 μM β-mercaptoethanol, 100 ng/mL human M-CSF (Peprotech), and 25 ng/mL human IL-3 (Peprotech). Every 7 days 2 ml media were replaced by fresh HM. After around 30 days, primitive macrophage precursors could be harvested during the media change and plated in microglia medium (MiM) at a density of 10^5^ cells/cm^2^. MiM consisted of Advanced DMEM F12 medium (Gibco) supplemented with 2 mM Glutamax, 100 U/mL penicillin, 100 μg/mL streptomycin, 55 μM β-mercaptoethanol, 100 ng/mL human IL-34 (Peprotech), and 10 ng/mL human GM-CSF (Peprotech). Finally, cells were differentiated in MiM for subsequent 6-9 days with full media change every other day.

### Generation of human hiPSC-derived cortical neurons

One day prior to neuronal induction, WTSIi015-A iPS cells were passaged using EDTA (Thermo Fisher Scientific) and pooled 2:1. The following day, medium was switched to neural maintenance media (NMM). NMM consisted of DMEM/F12 and neurobasal media (1:1) supplemented with 1x N2 supplement, 1x B27 supplement, 50 µM 2-mercaptoethanol, 0.5x non-essential amino acids, 100 µM L-glutamine (all from Life Technologies), 2500 U/mL penicillin/streptomycin (GE Healthcare), 10 µg/mL insulin and 0.5 mM sodium pyruvate (both from Sigma-Aldrich). NMM was further supplemented with 500 ng/mL mouse Noggin/CF chimera (R&D Systems) and 10 µM SB431542 (Stemcell Technologies). The cells were maintained in NMM for 10-12 days. The cells were then dissociated in colonies using 10 mg/ml Dispase II (Thermo Fisher Scientific) and seeded on laminin-coated plates (1-2 µg/cm^2^; Sigma-Aldrich) in NMM supplemented with 20 ng/mL FGF2 (Peprotech). The cells were kept in FGF2-supplemented medium for 4 to 5 days and then further passaged with dispase two times before day 25. After 25 days, the colonies were passaged and expanded using StemPro Accutase (Thermo Fisher Scientific) until day 35. Cells can be frozen between day 25-30. On day 35, the cells were passaged a last time onto plates coated with 1-2 µg/cm2 laminin at a density of 5*10^4^ cells/cm^2^ in NMM. The cells were then cultured for 2 weeks before they were used for experiments or the start of co-cultures with hiPSC-derived microglia.

### Co-culture of hiPSC-derived microglia and hiPSC-derived cortical neurons

Primitive macrophage precursors, harvested during media change, were directly added to hiPSC-derived cortical neurons at a density of 2.5*10^4^ cells/cm^2^. After 1 hour, media was changed to co-culture media (CoM). CoM consisted of DMEM/F12 and neurobasal media (1:1) supplemented with 1x N2 supplement, 2 mM Glutamax, 100 U/mL penicillin, 100 μg/mL streptomycin, 55 μM β-mercaptoethanol, 100 ng/mL human IL-34, and 10 ng/mL human GM-CSF. Cells were co-cultured for 2 weeks with full media change every second day.

### siRNA-mediated knockdown

Human hiPSC-derived microglia were transfected with Lipofectamine RNAiMAX reagent (Life Technologies) according to the manufactureŕs protocol. A final siRNA concentration of 200 nM was used. Cells were used for experiments on day 4 after transfection. The following Silencer siRNAs (Ambion) were used: control (siRNA ID: 4390843), TREM2 (siRNA ID: 289814) and cGAS (siRNA ID: 129127).

### Mice

C57BL/6 (WT) and *Trem2^-/-^* mice were bred at the University of Aarhus (Denmark). *Trem2^-/-^* mice were purchased from the Jackson Laboratory. Isoflurane (Abbott) or a mixture of ketamine (MSD Animal Health) and xylazine (Rompun Vet) were used to anesthetize mice. All described animal experiments have been reviewed and approved by the Danish government authorities and hence comply with Danish laws (ethics approval Nr. 2016−15−0201−01085). All efforts were made to minimize suffering and mice were monitored daily during infection. The mice were not randomized, but after HSV1 infection, the information about mouse strain and treatment was blinded to the investigators. No animals were excluded from the analysis. Chow and water were provided ad libitum.

### Murine ocular and HSV1 infection model

Age matched, 6–7-week-old male mice were anesthetized with i.p. injection of a mixture of ketamine (100 mg/kg body weight) and xylazine (10 mg/kg body weight). We tested male and female mice and did not find any sex differences in any of the readouts used in the current study. Both corneas were scarified in a 10 × 10 crosshatch pattern with a 25 gauge needle and mice inoculated with HSV1 (strain McKrae, the dosage used is indicated in the figure legends) in 5 μl of infection medium (DMEM containing 200 IU mL-1 penicillin and 200 mg mL-1 streptomycin), or mock-infected with 5 μl of infection medium.

### PFU assay

Vero cells (ATCC) were used for plaque assays and maintained in MEM containing 5% FBS, 1% penicillin, and 1% streptomycin. Vero cells were incubated with serial dilutions of cell supernatants. After 1 h, 0.75 % Methylcellulose (Sigma) was added, and plates were incubated at 37°C. After 3 days, the plaques were stained and counted under the microscope.

Mouse brainstems were isolated and immediately put on dry ice. Brainstems were then homogenized in DMEM and pelleted by centrifugation at 1600 g for 30 min. Supernatants were used for plaque assay as described (*12*).

### sTREM2 ELISA

sTREM2 was measured using an in-house electro-chemiluminescent assay on the MESO QuickPlex SQ 120 instrument (MesoScale Discovery, USA) using a method developed by Kleinberger et al. (*49*) with modifications by Alosco *et al.* (*50*). Briefly, the capture antibody was biotinylated polyclonal goat anti-human TREM2 (0.25 μg/mL, R&D Systems), and the detector antibody was monoclonal mouse anti-human TREM2 (1 μg/mL, Santa Cruz Biotechnology). A standard curve for calculations of unknowns was constructed using recombinant human TREM2 (4000–62.5 pg/mL, Sino Biological), and samples were diluted 1:4 before being assayed

### RNA isolation, RT-PCR, and qPCR

Tissues were homogenized with steel beads (Qiagen) in a Tissuelyser (II) (Qiagen) in PBS and immediately used for RNA isolation. RNA from mouse brainstem or primary cell cultures was isolated using the High Pure RNA Isolation Kit (Roche). HiPSC-derived cells were lysed in RLT Plus buffer (Qiagen) supplemented with 4 mM DTT (Sigma) and RNA was isolated using the RNeasy Mini Kit (Qiagen). cDNA was synthesized using the High-capacity cDNA Kit (Thermofisher).

Quantitative PCR was performed using the following TaqMan Gene Expression Assays (Applied Biosystems): *Actb* (Mm00607939_s1), *Trem2* (Mm04209424_g1), *Iba1* (Mm Mm00479862_g1), *ACTB* (Hs01060665_g1), *18S* (Hs03003631_g1), *TREM2* (Hs00219132_m1), *IFNB* (Hs01077958_s1), *IL6* (Hs00174131_m1), *TNFA* (Hs00174128_m1), *MX1* (Hs00895598_m1), and *cGAS* (Hs00403553_m1). For human *HSV1 gB* the following custom TaqMan primers and probe were used: forward primer 5′-GCAGTTTACGTACAACCACATACAGC-3′, reverse primer 5′-AGCTTGCGGGCCTCGTT-3′, and probe 56-FAM/CGGCCCAACATATCGTTGACATGGC/3BHQ_1 (IDT). For mouse *HSV1 gB* the following custom TaqMan primers and probe were used: forward primer (5′-CGCATCAAGACCACTCCTC-3′), reverse primer (5′-AGCTTGCGGGCCTCGTT-3′), and probe (5′-CGGCCCAACATATCGTTGACATGGC-3′). mRNA levels of interest were normalized to the housekeeping gene *Actb/ACTB* or *18S* (as indicated) using the ΔΔCT method. *18S* was used as a reference gene for co-cultures since *ACTB* expression was consistently affected by HSV1 infection.

### RNA sequencing and data analysis

HiPSC-derived microglia were grown on 24-well plates at a density of 2*10^5^ cells/well and infected with HSV1 (MOI1) for 24 h. Cells were then lysed in RLT Plus buffer supplemented with 4 mM DTT. RNA was isolated using the RNeasy Micro Kit (Qiagen) including the DNA digestion step and yielded ∼1.5 µg RNA. RNA-seq including bioinformatics analysis was performed at Omiics ApS (Denmark). Samples were rRNA depleted and prepared for sequencing using SMARTer Stranded Total RNA Sample Prep Kit - HI Mammalian (Takara). In brief, this kit first removes ribosomal RNA (rRNA) using RiboGone technology specifically depleting nuclear rRNA sequences (5S, 5.8S, 18S, and 28S) and mitochondrial rRNA 12S. RiboGone oligos are hybridized to rRNA. Which is cleaved using RNase H mediated cleavage. First strand synthesis is performed using random priming, adding an anchor for use with later PCR step. Template switching is utilized during the RT step and adds additional non-templated nucleotides to the 3’ end of the newly formed cDNA. PCR is performed leveraging the non-templated nucleotides and the added anchor sequence to produce Illumina compatible libraries. Prepared libraries were quality controlled using the Bioanalyzer 2100 (Agilent) and qPCR-based concentration measurements. Libraries were equimolarly pooled and sequenced including 150 bp paired end reads on an Illumina HiSeq sequencer. Sequencing data were pre-processed by removing adapter sequence and trimming away low-quality bases with a Phred1 score below 20 using Trim Galore (v0.4.1). Quality control was performed using FastQC, Picard and MultiQC to ensure high quality data. HSV1 genome (accession code JQ673480.1) was downloaded from NCBI. The filtered RNA-seq data was mapped against the viral genome using STAR. Reads not mapping to the viral genome were extracted and mapped against the human genome (hg38 / GRCh38) using STAR and gene expression was quantified using featureCounts with gene annotations from Gencode release 38. Differential expression analysis was performed using DESeq2 in R for human gene expression profiles.

Gene Ontology (GO) analysis was done using the clusterProfiler R package (*51*). For GO analysis all significantly differentially expressed genes (FDR < 0.05) were used except in cases were more than 3000 significant genes were found, in which case the 3000 most significant genes were used. Plotting was done in R.

### STRING analysis

To identify nodes in the virus-regulated transcripts, we used the protein-protein interaction analysis tool STRING (https://string-db.org/). The proteins corresponding to the 200 most downregulated genes in the RNA-seq dataset were subjected to STRING analysis. The analysis included only physical subnetworks and a confidence of at least 0.9 was used as cut-off criteria. At least three proteins were required to qualify for a node.

### Protein isolation and immunoblotting

HiPSC-derived microglia were grown on 6-well plates for protein isolation. Cells were washed 2x with PBS and lysed on ice for 5 min in RIPA buffer (20 mM Tris-HCl pH 7.5, 150 mM NaCl, 1 mM EDTA, 1% Triton X-100, 0.5% sodium deoxycholate, 0.1% SDS) supplemented with protease and phosphatase inhibitor cocktails (Roche). Samples were sonicated on ice for 10 min and centrifuged at 14000 g at 4°C for 10 min. Supernatants were collected and stored at -80°C for further use. For immunoblotting, samples were boiled at 95°C for 5 min under reducing conditions.

The following primary antibodies were used: TREM2 C-terminal (CS91068, Cell Signaling), GAPDH (CS5174), p-STING S366 (CS50907, Cell Signaling), t-STING (CS13647, Cell Signaling), p-TBK1 S172 (CS5483, Cell Signaling), t-TBK1 (CS3504, Cell Signaling), p-IRF3 S396 (CS29047, Cell Signaling), t-IRF3 (CS4302, Cell Signaling), cGAS (CS79978), beta-actin (CS3700, Cell Signaling), p-SYK Y525/526 (CS2710, Cell Signaling), t-SYK (CS13198, Cell Signaling). The following secondary antibodies were used: IRDye® 800CW donkey anti-rabbit IgG and IRDye® 680RD donkey anti-mouse IgG (both from LI-COR Biotechnology).

### Immunocytochemistry

Cells grown on Ibidi µ-slides (Ibidi) were washed 2x in PBS and fixed in 4% paraformaldehyde for 20 min at RT, covered with PBS and stored in fridge at 4°C until analysis. The cells were permeabilized with 0.3 % Triton-X100 in TBS for 15 min at RT and incubated with blocking buffer (0.3 % Triton-X100 and 5 % donkey serum in TBS) for 1 h at RT. Primary antibodies were diluted in blocking buffer. The following primary antibodies were used for staining: IBA1 (ab5076, Abcam), TUJ1 (ab14545, Abcam), and TREM2 (AF1828, R&D Systems). Samples were incubated with primary antibodies overnight at 4°C. Cells were washed 3x in TBS and incubated for 1 h with secondary antibodies diluted 1:500 in blocking buffer. Cells were washed 3x in TBS, counterstained using DAPI, and mounted using Ibidi mounting media (Ibidi). Samples were imaged using a Nikon A1 inverted confocal microscope.

### cGAMP quantification

HiPSC-derived microglia were grown on 6-well plates and infected with HSV1 (MOI 1) for 24 h. Cells were washed 2x with PBS, cells were carefully scraped, cells were collected in 500 µl PBS/well and 2 wells were merged. Cells were counted and then centrifuged at 200 g for 5 min. The supernatant was discarded and 250 µl internal standard solution (10 nM cGMP in acetonitrile:methanol [1:1]) was added to the cell pellet. Samples were vortexed and stored at -80°C until further analysis. Samples were extracted by sonication for 10 min followed by vortexing (1400 rpm) for an additional 10 min. Samples were centrifuged at 16000 g for 10 min and supernatants were evaporated under a stream of nitrogen before reconstitution in 100 µl acetonitrile:methanol [3:1]. Quantification was performed using ultra-performance liquid chromatography coupled to tandem mass spectrometry (UPLC-MS/MS). Five µl were injected onto a Waters zHILIC column (2.1×100; 1.7 µM particles) and separated using a gradient of acetonitrile with 0.1% formic acid as B-phase and water with 20 mM ammonium formate as A-phase. Detection was performed on a Sciex 7500 triple quadrupole mass spectrometer (Sciex). Quantification was made against an external calibration curve with a limit of detection of 0.1nM.

### Cell viability assay

The CyQUANT XTT assay (Invitrogen) was used to measure cell viability. HiPSC-derived co-cultures were infected with HSV1 as indicated. After 24 h, 150 µl media were removed and stored at -80°C for further analyses. Sixty-five µl CyQUANT Mix were added to the remaining 100 µl and cells were incubated for 3 h. Cell supernatants were transferred to a 96-well plate and absorbance (A) was measured at 450 and 660 nm. Specific absorbance was calculated by using the formula: A_(450 nm)_-A_(660 nm)_.

### Annexin V staining and FACS analysis

Annexin V-Alexa 488 conjugates (Thermofisher) were used for apoptosis detection according to the manufactureŕs instructions. Briefly, hiPSC-derived microglia were trypsinized, washed 1x with PBS at 400 g for 5 min, and resuspended in annexin-binding buffer at 1*10^6^ cells/ml. 5 µl annexin V conjugate were added per 100 µl cell solution, incubated for 15 min at RT, and 400 µl of annexin-binding buffer were added. Cells were analyzed using a FACSAria Fusion, the Annexin V conjugate signal was detected in the FITC channel and positive cells were quantified using the BD FACSDiva software.

### Phagocytosis assays

For analysis of phagocytosis of HSV1-infected neurons, hiPSC-derived neurons were infected with HSV1 (10 MOI). After 24 h, cells were washed 1x with PBS and fresh NMM was added. Cells were subjected to UV irradiation for 10 min to inactivate HSV1. Cells were then trypsinized, centrifuged at 400 g for 5 min, resuspended in PBS containing 2 µg/ml fluorescent cell tracker (CellTracker™ CM-DiI Dye, Thermofisher), and incubated for 5 min at 37°C. Cells were washed 2x with PBS at 400 g for 5 min. The UV inactivation of the virus was confirmed by plaque assay. Cells were resuspended in MiM and added to the cultured hiPSC-derived microglia, at a 1:1 cell ratio, for 15 h. For ICC, microglia were washed 2x with PBS and fixed in 4% paraformaldehyde for 20 min at RT followed by DAPI counterstaining and mounting. Samples were imaged using a Nikon A1 inverted confocal microscope. For FACS analysis, microglia were trypsinized, washed 2x with PBS at 400 g for 5 min, and resuspended in 200 µl PBS for FACS analysis using a FACSAria Fusion. The cell tracker signal was detected in the PE channel and positive cells were quantified using the BD FACSDiva software.

For analysis of phagocytosis of apoptotic neurons, apoptosis was induced in hiPSC-derived neurons by adding 30 nM okadaic acid to the cell media for 3 h. Neurons were trypsinized, washed 1x in PBS, resuspended in PBS containing 2 µg/ml fluorescent cell tracker, and incubated for 5 min at 37°C. Cells were washed 2x with PBS at 400 g for 5 min, resuspended in MiM, and added to the cultured hiPSC-derived microglia, at a 1:10 cell ratio, for 3 h. Microglia were prepared for FACS analysis as above.

For analysis of phagocytosis of pHrodo-*E. coli* beads (Thermofisher), beads were resuspended in live cell imaging solution (2 ml/vial of beads, Thermofisher) and briefly sonicated. hiPSC-derived microglia were grown in black clear bottom 96-well plates and incubated with 100 µl beads solution for 3 h. Phagocytosis of pHrodo-*E. coli* beads was quantified using a microplate reader and excitation/emission wavelengths of 560/585 nm.

To inhibit phagocytosis, cells were incubated with 5 µM Cytochalasin D (Thermofisher) 1 hour prior to experiments and during experiments.

### Statistical analyses

For statistical analysis of data, we used 2-tailed Student’s t test when the data exhibited normal distribution, and Mann Whitney U test when the data set did not pass the normal distribution test or normal distribution could not be tested. When comparing more than 2 groups, multiple-comparison 1-way ANOVA was used with Tukey’s multiple-comparison test. All experimental data were reliably reproduced in 2 or more independent experiments. Individual data points represent biological replicates. GraphPad Prism 8 software and R were used for statistical analyses. No measurement was excluded for statistical analysis.

## List of Supplementary Materials

**Fig. S1.**
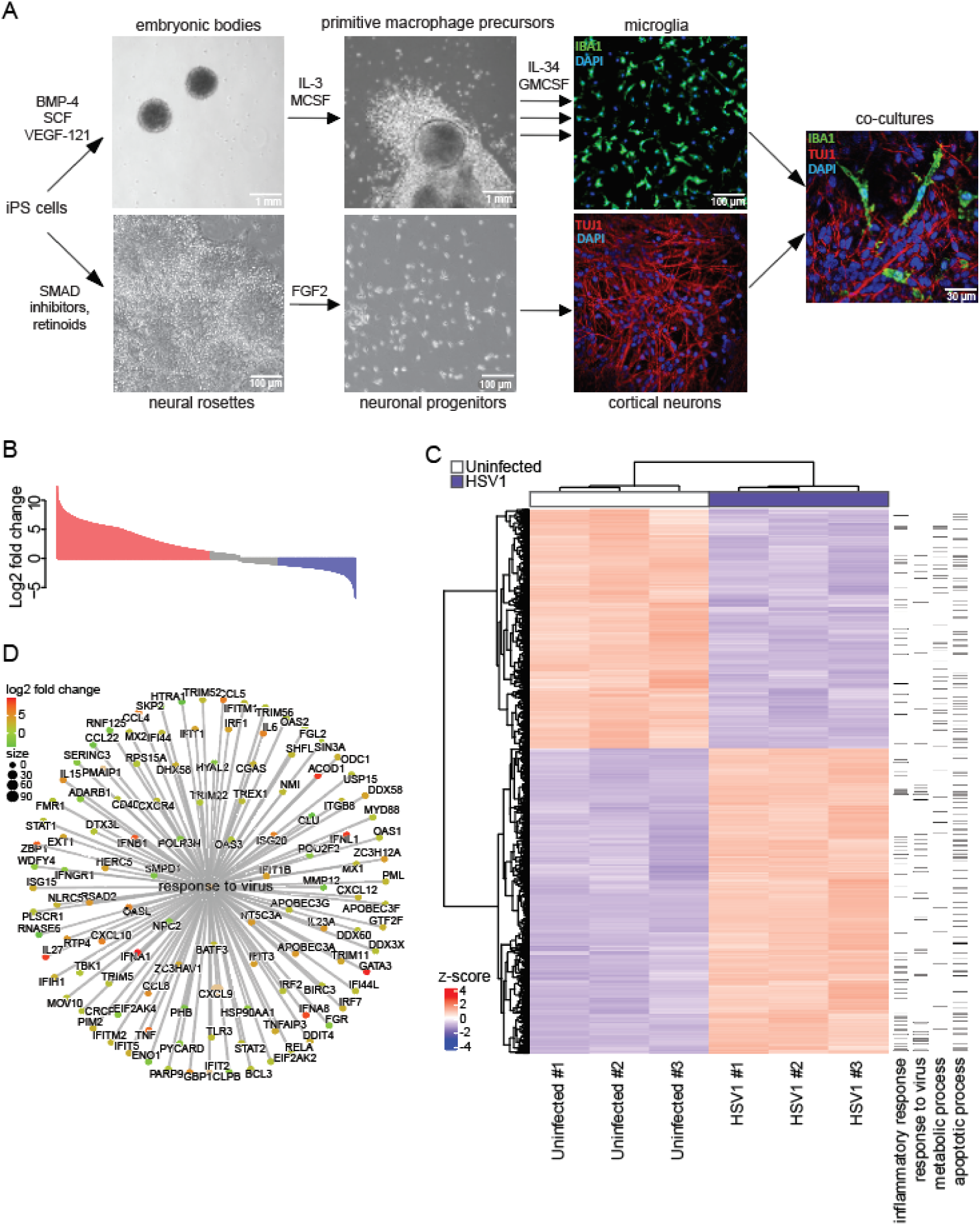
Stem cell differentiation workflow and RNA-seq data analysis. (**A**) Illustration of the differentiation workflow from hiPSCs towards microglia and cortical neurons is shown. Cells were stained for markers for microglia (IBA1, green), cortical neurons (TUJ1, red), and nuclei (DAPI, blue). Multiple arrows indicate continuous harvest of microglia. (**B**) Waterfall plot for RNA-seq of HSV1-infected microglia relative to mock infection. (**C**) Hierarchical clustering of genes differentially expressed in uninfected versus HSV1-infected microglia. To the right biological function annotated with the regulated genes is shown. (**D**) Differentially regulated genes in the gene ontology category “response to virus” in HSV-infected microglia.

**Fig. S2.**
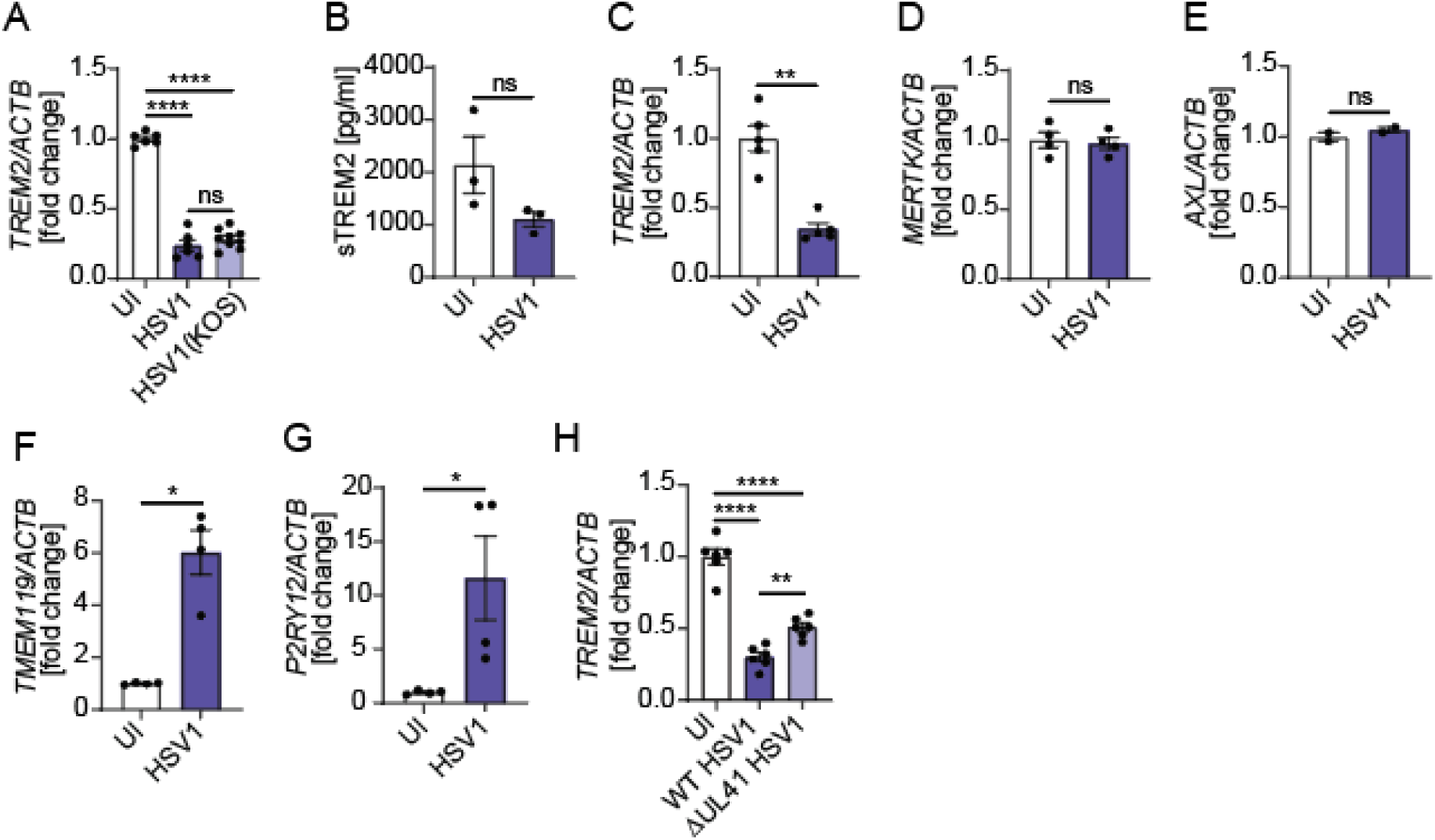
HSV1 downregulates microglial TREM2 expression independent of the viral strain and iPSC line used. (**A**) *TREM2* mRNA levels were analyzed 24 h post-infection of hiPSC-derived microglia with the clinical HSV1 strain (MOI 1) and the KOS strain (MOI 1). (**B**-**C**) Microglia differentiated from a separate iPSC line (ChiPSC22). (**B**) sTREM2 levels were analyzed 24 h post-infection with HSV1 (MOI 1) by ELISA. (**C**) *TREM2* mRNA levels were analyzed 24 h post-infection with HSV1 (MOI 1). (**D**-**G**) Examples of genes which mRNA levels were unchanged (*MERTK* and *AXL*) or upregulated (*TMEM119* and *P2RY12*) in microglia 24 h post-infection with HSV1 (MOI 1). (**H**) *TREM2* mRNA levels were analyzed 24 h post-infection with wildtype (WT) HSV1 and a UL41 mutant strain (ΔUL41). All figures represent 2-3 independent experiments; data are presented as mean ± SEM; P values were calculated by one-way ANOVA with Tukey’s multiple comparisons test (A and H) and Mann-Whitney test (B-G). * P < 0.05; ** P < 0.001; **** P < 0.0001.

**Fig. S3.**
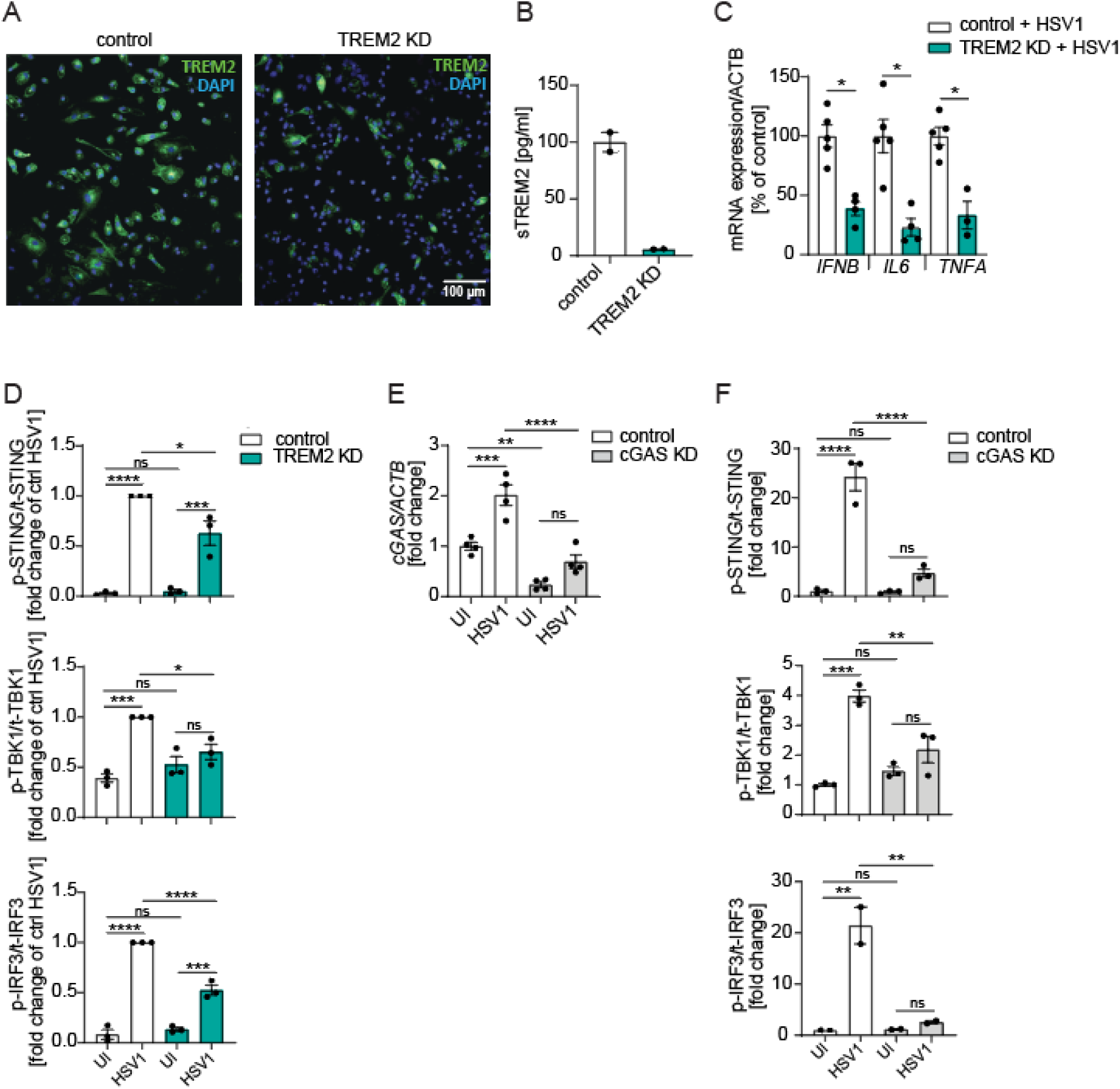
Characterization of TREM2 KD hiPSC-derived microglia. (**A**) Control and TREM2 KD microglia were stained for TREM2 (green) and DAPI (blue) and imaged using a confocal microscope. (**B**) sTREM2 levels were measured in in cell supernatants by ELISA. All figures represent 2 independent experiments; data are presented as mean ± SEM. (**C**) Microglia differentiated from a separate iPSC line (ChiPSC22). *IFNB*, *IL6*, and *TNFA* mRNA levels were analyzed 24 h post-infection with HSV1 (MOI 3). (**D**) Quantifications of Fig. 3F (**E**) cGAS was knocked down in microglia using siRNA. Cells were used for experiments 4 days after transfection. Control and cGAS KD microglia were infected with HSV1 (MOI 3) for 24 h. *cGAS* mRNA levels were quantified. (**F**) Quantifications of Fig. 2K. All figures represent 2-3 independent experiments; data are presented as mean ± SEM; P values were calculated by Mann-Whitney test (B-C) and one-way ANOVA with Tukey’s multiple comparisons test (D-F). * P < 0.05; ** P < 0.001; *** P < 0.0005; **** P< 0.0001.

**Fig. S4.**
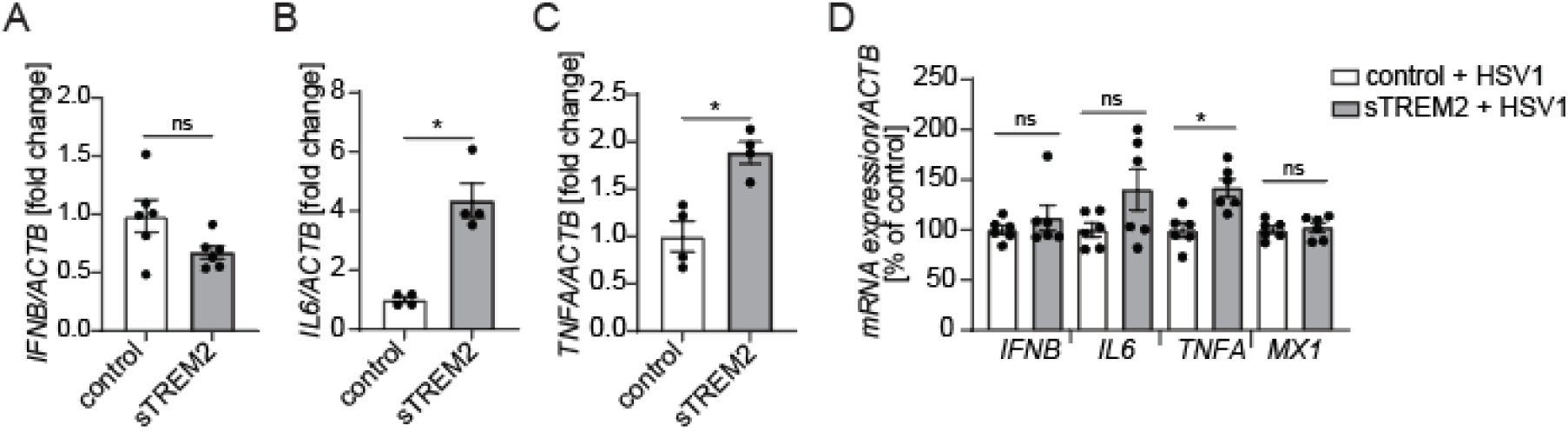
sTREM2 activates inflammatory responses but not antiviral defense. (**A**-**C**) HiPSC-derived microglia were incubated with 3 µg/ml sTREM2 for 24 h. *IFNB*, *IL6*, and *TNFA* mRNA levels are shown. (**D**) *IFNB*, *IL6*, *TNFA*, and *MX1* mRNA levels were analyzed in microglia after infection with HSV1 (MOI 1) with and without sTREM2 for 8 h. All figures represent 2-3 independent experiments; data are presented as mean ± SEM; P values were calculated by Mann-Whitney test. * P < 0.05.

**Fig. S5.**
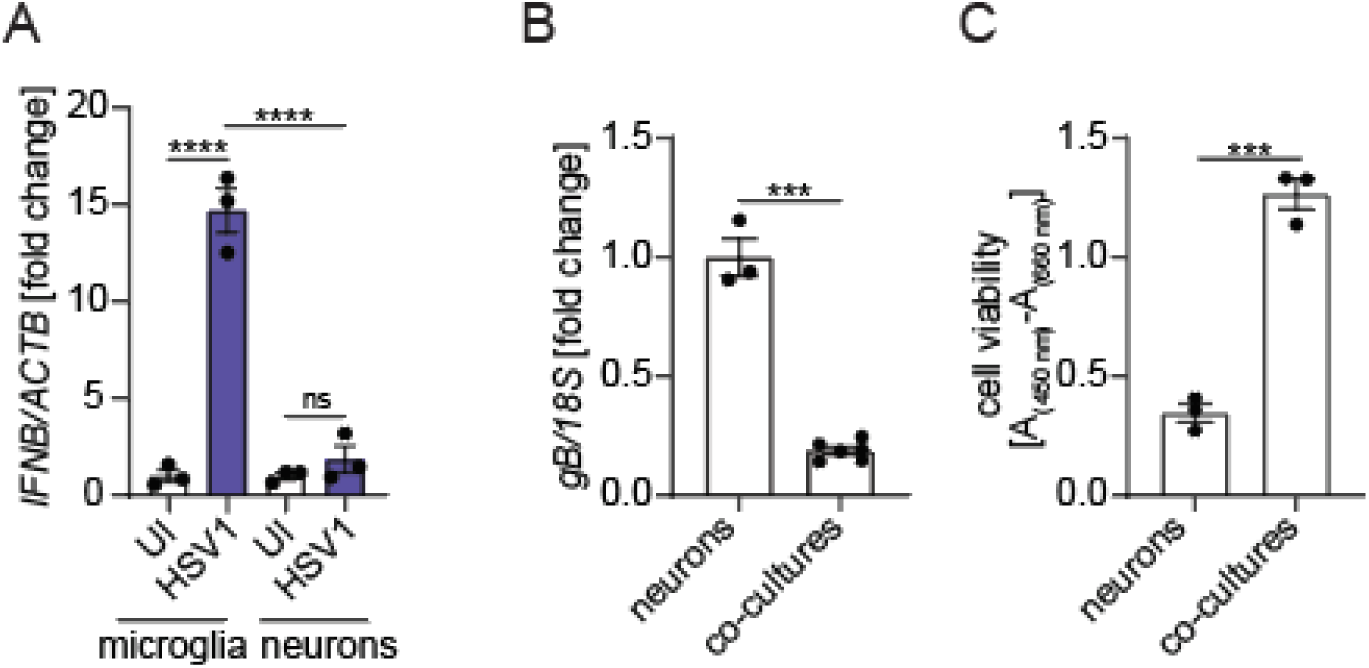
Microglia are the main source of IFNB and decrease viral replication in co-cultures compared with neurons alone. (**A**) *IFNB* mRNA levels were analyzed in hiPSC-derived microglia and neurons 24 h after infection with HSV1 (MOI 1). (**B**, **C**) HiPSC-derived neurons and co-cultures of microglia and neurons were analyzed 24 h after infection with HSV1 (MOI 1). (**B**) *gB* mRNA levels are shown. (**C**) Cell viability is shown. All figures represent 1-2 independent experiments; data are presented as mean ± SEM; P values were calculated by two-way ANOVA with Tukey’s multiple comparisons test (A) and 2-tailed Student’s t test (B, C). *** P < 0.0005; **** P < 0.0001.

**Fig. S6.**
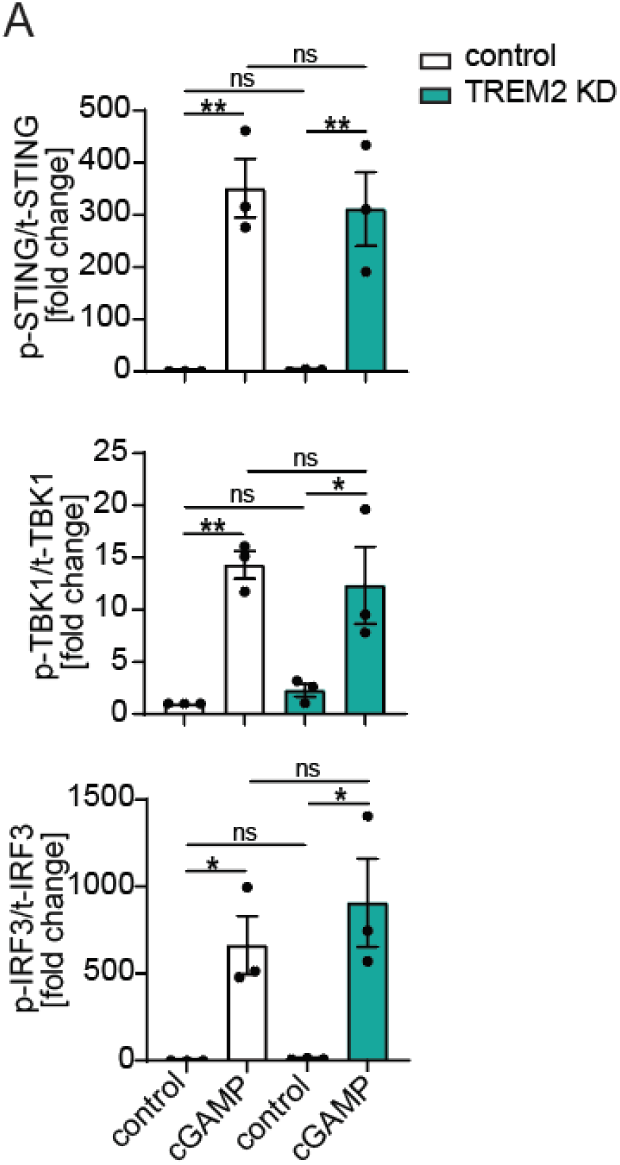
Quantification of cGAS/STING signaling in cGAMP-stimulated microglia. (**A**) Quantifications of Fig.5B are shown. All figures represent 3 independent experiments; data are presented as mean ± SEM; P values were calculated by one-way ANOVA with Tukey’s multiple comparisons test. * P < 0.05; ** P < 0.001.

Table S1 to S2

## Acknowledgments

We would like to thank Monica Malmberg for her help with sTREM2 immunossays and Susann Li for her help with FACS. We would like to acknowledge that Fig. 6E was created with BioRender.com.

## Funding

Petrus och Augusta Hedlunds Stiftelse M-2022-1825 (SF)

The European Research Council 786602 (SRP)

The Lundbeck Foundation R359-2020-2287 (SRP)

The Swedish Research Council 2018-02463 and 2021-00942 (SRP)

The Novo Nordisk Foundation NNF20OC0063436 (SRP)

Swedish Research Council 2018-02532 (HZ)

European Research Council 681712 (HZ)

European Research Council 101053962 (HZ)

Swedish State Support for Clinical Research ALFGBG-71320 (HZ)

Alzheimer Drug Discovery Foundation 201809-2016862 (HZ)

Alzheimer’s Association ADSF-21-831376-C (HZ)

Alzheimer’s Association ADSF-21-831381-C (HZ)

Alzheimer’s Association ADSF-21-831377-C (HZ)

Hjärnfonden FO2019-0228 (HZ)

European Union’s Horizon 2020 research and innovation programme under the Marie Skłodowska-Curie grant agreement 860197 (HZ)

European Union Joint Programme-Neurodegenerative Disease Research JPND2021-00694 (HZ)

UK Dementia Research Institute at UCL UKDRI-1003 (HZ)

## Author contributions

Conceptualization: SF, SRP, LSR, HZ

Methodology: SF, LSR, MH

Investigation: SF, LSR, CÖ, MS, MH

Visualization: SF

Funding acquisition: SRP, HZ, SF

Supervision: SRP, HZ

Writing – original draft: SF, SRP

Writing – review & editing: SF, SRP, HZ, LSR, CÖ, MS, MH

## Data and materials availability

All data are available in the main text or the supplementary materials.

## References

1. K. L. Tyler, Acute Viral Encephalitis. N Engl J Med 379, 557–566 (2018).

2. J. P. Stahl, A. Mailles, Herpes simplex virus encephalitis update. Curr Opin Infect Dis 32, 239–243 (2019).

2. K. L. Roos, Encephalitis. Handb Clin Neurol 121, 1377–1381 (2014).

3. R. G. Pebody, N. Andrews, D. Brown, R. Gopal, H. De Melker, G. Francois, N. Gatcheva, W. Hellenbrand, S. Jokinen, I. Klavs, M. Kojouharova, T. Kortbeek, B. Kriz, K. Prosenc, K. Roubalova, P. Teocharov, W. Thierfelder, M. Valle, P. Van Damme, R. Vranckx, The seroepidemiology of herpes simplex virus type 1 and 2 in Europe. Sex Transm Infect 80, 185–191 (2004).

4. A. J. Steel, G. D. Eslick, Herpes Viruses Increase the Risk of Alzheimer’s Disease: A Meta-Analysis. Journal of Alzheimer’s disease : JAD 47, 351–364 (2015).

5. R. F. Itzhaki, W. R. Lin, D. Shang, G. K. Wilcock, B. Faragher, G. A. Jamieson, Herpes simplex virus type 1 in brain and risk of Alzheimer’s disease. Lancet 349, 241–244 (1997).

6. O. Uyar, N. Laflamme, J. Piret, M.-C. Venable, J. Carbonneau, K. Zarrouk, S. Rivest, G. Boivin, An Early Microglial Response Is Needed To Efficiently Control Herpes Simplex Virus Encephalitis. Journal of Virology 94, (2020).

7. R. Fekete, C. Cserep, N. Lenart, K. Toth, B. Orsolits, B. Martinecz, E. Mehes, B. Szabo, V. Nemeth, B. Gonci, B. Sperlagh, Z. Boldogkoi, A. Kittel, M. Baranyi, S. Ferenczi, K. Kovacs, G. Szalay, B. Rozsa, C. Webb, G. G. Kovacs, T. Hortobagyi, B. L. West, Z. Kornyei, A. Denes, Microglia control the spread of neurotropic virus infection via P2Y12 signalling and recruit monocytes through P2Y12-independent mechanisms. Acta Neuropathol 136, 461–482 (2018).

8. G. Katzilieris-Petras, X. Lai, A. S. Rashidi, G. Verjans, L. S. Reinert, S. R. Paludan, Microglia activate early anti-viral responses upon HSV-1 entry into the brain to counteract development of encephalitis-like disease in mice. *J Virol*, JVI0131121 (2022).

9. L. Sun, J. Wu, F. Du, X. Chen, Z. J. Chen, Cyclic GMP-AMP synthase is a cytosolic DNA sensor that activates the type I interferon pathway. Science 339, 786–791 (2013).

10. H. Ishikawa, G. N. Barber, STING is an endoplasmic reticulum adaptor that facilitates innate immune signalling. Nature 455, 674–678 (2008).

11. L. S. Reinert, K. Lopusna, H. Winther, C. Sun, M. K. Thomsen, R. Nandakumar, T. H. Mogensen, M. Meyer, C. Vaegter, J. R. Nyengaard, K. A. Fitzgerald, S. R. Paludan, Sensing of HSV-1 by the cGAS-STING pathway in microglia orchestrates antiviral defence in the CNS. Nat Commun 7, 13348 (2016).

12. M. Herman, M. Ciancanelli, Y. H. Ou, L. Lorenzo, M. Klaudel-Dreszler, E. Pauwels, V. Sancho-Shimizu, R. Perez de Diego, A. Abhyankar, E. Israelsson, Y. Guo, A. Cardon, F. Rozenberg, P. Lebon, M. Tardieu, E. Heropolitanska-Pliszka, D. Chaussabel, M. A. White, L. Abel, S. Y. Zhang, J. L. Casanova, Heterozygous TBK1 mutations impair TLR3 immunity and underlie herpes simplex encephalitis of childhood. J Exp Med 209, 1567–1582 (2012).

13. L. L. Andersen, N. Mork, L. S. Reinert, E. Kofod-Olsen, R. Narita, S. E. Jorgensen, K. A. Skipper, K. Honing, H. H. Gad, L. Ostergaard, T. F. Orntoft, V. Hornung, S. R. Paludan, J. G. Mikkelsen, T. Fujita, M. Christiansen, R. Hartmann, T. H. Mogensen, Functional IRF3 deficiency in a patient with herpes simplex encephalitis. J Exp Med 212, 1371–1379 (2015).

14. S. Pan, X. Liu, Y. Ma, Y. Cao, B. He, Herpes Simplex Virus 1 gamma134.5 Protein Inhibits STING Activation That Restricts Viral Replication. J Virol 92, (2018).

15. J. Huang, H. You, C. Su, Y. Li, S. Chen, C. Zheng, Herpes Simplex Virus 1 Tegument Protein VP22 Abrogates cGAS/STING-Mediated Antiviral Innate Immunity. J Virol 92, (2018).

16. M. H. Christensen, S. B. Jensen, J. J. Miettinen, S. Luecke, T. Prabakaran, L. S. Reinert, T. Mettenleiter, Z. J. Chen, D. M. Knipe, R. M. Sandri-Goldin, L. W. Enquist, R. Hartmann, T. H. Mogensen, S. A. Rice, T. A. Nyman, S. Matikainen, S. R. Paludan, HSV-1 ICP27 targets the TBK1-activated STING signalsome to inhibit virus-induced type I IFN expression. EMBO J 35, 1385–1399 (2016).

17. C. Bodda, L. S. Reinert, S. Fruhwürth, T. Richardo, C. Sun, B. C. Zhang, M. Kalamvoki, A. Pohlmann, T. H. Mogensen, P. Bergström, L. Agholme, P. O’Hare, B. Sodeik, M. Gyrd-Hansen, H. Zetterberg, S. R. Paludan, HSV1 VP1-2 deubiquitinates STING to block type I interferon expression and promote brain infection. J Exp Med 217, (2020).

18. D. L. Kober, T. J. Brett, TREM2-Ligand Interactions in Health and Disease. J Mol Biol 429, 1607–1629 (2017).

19. F. Filipello, C. Goldsbury, S. F. You, A. Locca, C. M. Karch, L. Piccio, Soluble TREM2: Innocent bystander or active player in neurological diseases? Neurobiology of disease 165, 105630 (2022).

20. J. Paloneva, M. Kestila, J. Wu, A. Salminen, T. Bohling, V. Ruotsalainen, P. Hakola, A. B. Bakker, J. H. Phillips, P. Pekkarinen, L. L. Lanier, T. Timonen, L. Peltonen, Loss-of-function mutations in TYROBP (DAP12) result in a presenile dementia with bone cysts. Nat Genet 25, 357–361 (2000).

21. J. Paloneva, T. Manninen, G. Christman, K. Hovanes, J. Mandelin, R. Adolfsson, M. Bianchin, T. Bird, R. Miranda, A. Salmaggi, L. Tranebjaerg, Y. Konttinen, L. Peltonen, Mutations in two genes encoding different subunits of a receptor signaling complex result in an identical disease phenotype. Am J Hum Genet 71, 656–662 (2002).

22. R. Guerreiro, A. Wojtas, J. Bras, M. Carrasquillo, E. Rogaeva, E. Majounie, C. Cruchaga, C. Sassi, J. S. Kauwe, S. Younkin, L. Hazrati, J. Collinge, J. Pocock, T. Lashley, J. Williams, J. C. Lambert, P. Amouyel, A. Goate, R. Rademakers, K. Morgan, J. Powell, P. St George-Hyslop, A. Singleton, J. Hardy, G. Alzheimer Genetic Analysis, TREM2 variants in Alzheimer’s disease. N Engl J Med 368, 117–127 (2013).

23. T. Jonsson, H. Stefansson, S. Steinberg, I. Jonsdottir, P. V. Jonsson, J. Snaedal, S. Bjornsson, J. Huttenlocher, A. I. Levey, J. J. Lah, D. Rujescu, H. Hampel, I. Giegling, O. A. Andreassen, K. Engedal, I. Ulstein, S. Djurovic, C. Ibrahim-Verbaas, A. Hofman, M. A. Ikram, C. M. van Duijn, U. Thorsteinsdottir, A. Kong, K. Stefansson, Variant of TREM2 associated with the risk of Alzheimer’s disease. N Engl J Med 368, 107–116 (2013).

24. G. Katzilieris-Petras, X. Lai, A. S. Rashidi, G. Verjans, L. S. Reinert, S. R. Paludan, Microglia Activate Early Antiviral Responses upon Herpes Simplex Virus 1 Entry into the Brain to Counteract Development of Encephalitis-Like Disease in Mice. J Virol 96, e0131121 (2022).

25. L. S. Reinert, A. S. Rashidi, D. N. Tran, G. Katzilieris-Petras, A. K. Hvidt, M. Gohr, S. Fruhwürth, C. Bodda, M. K. Thomsen, M. H. Vendelbo, A. R. Khan, B. Hansen, P. Bergström, L. Agholme, T. H. Mogensen, M. H. Christensen, J. R. Nyengaard, G. C. Sen, H. Zetterberg, G. M. Verjans, S. R. Paludan, Brain immune cells undergo cGAS/STING-dependent apoptosis during herpes simplex virus type 1 infection to limit type I IFN production. The Journal of clinical investigation 131, (2021).

26. G. C. Brown, P. St George-Hyslop, Does Soluble TREM2 Protect Against Alzheimer’s Disease? Front Aging Neurosci 13, 834697 (2021).

27. L. Zhong, Y. Xu, R. Zhuo, T. Wang, K. Wang, R. Huang, D. Wang, Y. Gao, Y. Zhu, X. Sheng, K. Chen, N. Wang, L. Zhu, D. Can, Y. Marten, M. Shinohara, C. C. Liu, D. Du, H. Sun, L. Wen, H. Xu, G. Bu, X. F. Chen, Soluble TREM2 ameliorates pathological phenotypes by modulating microglial functions in an Alzheimer’s disease model. Nat Commun 10, 1365 (2019).

28. P. Garcia-Reitboeck, A. Phillips, T. M. Piers, C. Villegas-Llerena, M. Butler, A. Mallach, C. Rodrigues, C. E. Arber, A. Heslegrave, H. Zetterberg, H. Neumann, S. Neame, H. Houlden, J. Hardy, J. M. Pocock, Human Induced Pluripotent Stem Cell-Derived Microglia-Like Cells Harboring TREM2 Missense Mutations Show Specific Deficits in Phagocytosis. Cell Rep 24, 2300–2311 (2018).

29. C. Wang, N. Sharma, M. Veleeparambil, P. M. Kessler, B. Willard, G. C. Sen, STING-Mediated Interferon Induction by Herpes Simplex Virus 1 Requires the Protein Tyrosine Kinase Syk. mBio 12, e0322821 (2021).

30. A. McQuade, Y. J. Kang, J. Hasselmann, A. Jairaman, A. Sotelo, M. Coburn, S. K. Shabestari, J. P. Chadarevian, G. Fote, C. H. Tu, E. Danhash, J. Silva, E. Martinez, C. Cotman, G. A. Prieto, L. M. Thompson, J. S. Steffan, I. Smith, H. Davtyan, M. Cahalan, H. Cho, M. Blurton-Jones, Gene expression and functional deficits underlie TREM2-knockout microglia responses in human models of Alzheimer’s disease. Nat Commun 11, 5370 (2020).

31. C. Kim, E. D. Cho, H. K. Kim, S. You, H. J. Lee, D. Hwang, S. J. Lee, beta1-integrin-dependent migration of microglia in response to neuron-released alpha-synuclein. Exp Mol Med 46, e91 (2014).

32. K. Takahashi, C. D. Rochford, H. Neumann, Clearance of apoptotic neurons without inflammation by microglial triggering receptor expressed on myeloid cells-2. J Exp Med 201, 647–657 (2005).

33. P. Garcia-Reitboeck, A. Phillips, T. M. Piers, C. Villegas-Llerena, M. Butler, A. Mallach, C. Rodrigues, C. E. Arber, A. Heslegrave, H. Zetterberg, H. Neumann, S. Neame, H. Houlden, J. Hardy, J. M. Pocock, Human Induced Pluripotent Stem Cell-Derived Microglia-Like Cells Harboring TREM2 Missense Mutations Show Specific Deficits in Phagocytosis. Cell Reports 24, 2300–2311 (2018).

34. C. L. Hsieh, M. Koike, S. C. Spusta, E. C. Niemi, M. Yenari, M. C. Nakamura, W. E. Seaman, A role for TREM2 ligands in the phagocytosis of apoptotic neuronal cells by microglia. J Neurochem 109, 1144–1156 (2009).

35. L. S. Reinert, A. S. Rashidi, D. N. Tran, G. Katzilieris-Petras, A. K. Hvidt, M. Gohr, S. Fruhwürth, C. Bodda, M. K. Thomsen, M. H. Vendelbo, A. R. Khan, B. Hansen, P. Bergström, L. Agholme, T. H. Mogensen, M. H. Christensen, J. R. Nyengaard, G. C. Sen, H. Zetterberg, G. M. Verjans, S. R. Paludan, Brain immune cells undergo cGAS/STING-dependent apoptosis during herpes simplex virus type 1 infection to limit type I IFN production. Journal of Clinical Investigation 131, (2021).

36. F. Mazaheri, N. Snaidero, G. Kleinberger, C. Madore, A. Daria, G. Werner, S. Krasemann, A. Capell, D. Trumbach, W. Wurst, B. Brunner, S. Bultmann, S. Tahirovic, M. Kerschensteiner, T. Misgeld, O. Butovsky, C. Haass, TREM2 deficiency impairs chemotaxis and microglial responses to neuronal injury. EMBO Rep 18, 1186–1198 (2017).

37. S. Liu, Y. Liao, B. Chen, Y. Chen, Z. Yu, H. Wei, L. Zhang, S. Huang, P. B. Rothman, G. F. Gao, J. L. Chen, Critical role of Syk-dependent STAT1 activation in innate antiviral immunity. Cell Rep 34, 108627 (2021).

38. L. N. Zhang, M. J. Li, Y. H. Shang, F. F. Zhao, H. C. Huang, F. X. Lao, Independent and Correlated Role of Apolipoprotein E varepsilon4 Genotype and Herpes Simplex Virus Type 1 in Alzheimer’s Disease. Journal of Alzheimer’s disease : JAD 77, 15–31 (2020).

39. 2021 Alzheimer’s disease facts and figures. Alzheimer’s & dementia : the journal of the Alzheimer’s Association 17, 327–406 (2021).

40. Y. Shi, D. M. Holtzman, Interplay between innate immunity and Alzheimer disease: APOE and TREM2 in the spotlight. Nat Rev Immunol 18, 759–772 (2018).

41. H. Keren-Shaul, A. Spinrad, A. Weiner, O. Matcovitch-Natan, R. Dvir-Szternfeld, T. K. Ulland, E. David, K. Baruch, D. Lara-Astaiso, B. Toth, S. Itzkovitz, M. Colonna, M. Schwartz, I. Amit, A Unique Microglia Type Associated with Restricting Development of Alzheimer’s Disease. Cell 169, 1276–1290 e1217 (2017).

42. Y. Wang, T. K. Ulland, J. D. Ulrich, W. Song, J. A. Tzaferis, J. T. Hole, P. Yuan, T. E. Mahan, Y. Shi, S. Gilfillan, M. Cella, J. Grutzendler, R. B. DeMattos, J. R. Cirrito, D. M. Holtzman, M. Colonna, TREM2-mediated early microglial response limits diffusion and toxicity of amyloid plaques. J Exp Med 213, 667–675 (2016).

43. S. Parhizkar, T. Arzberger, M. Brendel, G. Kleinberger, M. Deussing, C. Focke, B. Nuscher, M. Xiong, A. Ghasemigharagoz, N. Katzmarski, S. Krasemann, S. F. Lichtenthaler, S. A. Muller, A. Colombo, L. S. Monasor, S. Tahirovic, J. Herms, M. Willem, N. Pettkus, O. Butovsky, P. Bartenstein, D. Edbauer, A. Rominger, A. Erturk, S. A. Grathwohl, J. J. Neher, D. M. Holtzman, M. Meyer-Luehmann, C. Haass, Loss of TREM2 function increases amyloid seeding but reduces plaque-associated ApoE. Nat Neurosci 22, 191–204 (2019).

44. S. Wang, M. Mustafa, C. M. Yuede, S. V. Salazar, P. Kong, H. Long, M. Ward, O. Siddiqui, R. Paul, S. Gilfillan, A. Ibrahim, H. Rhinn, I. Tassi, A. Rosenthal, T. Schwabe, M. Colonna, Anti-human TREM2 induces microglia proliferation and reduces pathology in an Alzheimer’s disease model. J Exp Med 217, (2020).

45. K. Schlepckow, K. M. Monroe, G. Kleinberger, L. Cantuti-Castelvetri, S. Parhizkar, D. Xia, M. Willem, G. Werner, N. Pettkus, B. Brunner, A. Sulzen, B. Nuscher, H. Hampel, X. Xiang, R. Feederle, S. Tahirovic, J. I. Park, R. Prorok, C. Mahon, C. C. Liang, J. Shi, D. J. Kim, H. Sabelstrom, F. Huang, G. Di Paolo, M. Simons, J. W. Lewcock, C. Haass, Enhancing protective microglial activities with a dual function TREM2 antibody to the stalk region. EMBO Mol Med 12, e11227 (2020).

46. J. Piret, G. Boivin, Immunomodulatory Strategies in Herpes Simplex Virus Encephalitis. Clinical Microbiology Reviews 33, (2020).

47. B. Skoldenberg, M. Forsgren, K. Alestig, T. Bergstrom, L. Burman, E. Dahlqvist, A. Forkman, A. Fryden, K. Lovgren, K. Norlin, et al., Acyclovir versus vidarabine in herpes simplex encephalitis. Randomised multicentre study in consecutive Swedish patients. Lancet 2, 707–711 (1984).

48. G. Kleinberger, Y. Yamanishi, M. Suarez-Calvet, E. Czirr, E. Lohmann, E. Cuyvers, H. Struyfs, N. Pettkus, A. Wenninger-Weinzierl, F. Mazaheri, S. Tahirovic, A. Lleo, D. Alcolea, J. Fortea, M. Willem, S. Lammich, J. L. Molinuevo, R. Sanchez-Valle, A. Antonell, A. Ramirez, M. T. Heneka, K. Sleegers, J. van der Zee, J. J. Martin, S. Engelborghs, A. Demirtas-Tatlidede, H. Zetterberg, C. Van Broeckhoven, H. Gurvit, T. Wyss-Coray, J. Hardy, M. Colonna, C. Haass, TREM2 mutations implicated in neurodegeneration impair cell surface transport and phagocytosis. Sci Transl Med 6, 243ra286 (2014).

49. M. L. Alosco, Y. Tripodis, N. G. Fritts, A. Heslegrave, C. M. Baugh, S. Conneely, M. Mariani, B. M. Martin, S. Frank, J. Mez, T. D. Stein, R. C. Cantu, A. C. McKee, L. M. Shaw, J. Q. Trojanowski, K. Blennow, H. Zetterberg, R. A. Stern, Cerebrospinal fluid tau, Abeta, and sTREM2 in Former National Football League Players: Modeling the relationship between repetitive head impacts, microglial activation, and neurodegeneration. Alzheimer’s & dementia : the journal of the Alzheimer’s Association 14, 1159–1170 (2018).

50. T. Wu, E. Hu, S. Xu, M. Chen, P. Guo, Z. Dai, T. Feng, L. Zhou, W. Tang, L. Zhan, X. Fu, S. Liu, X. Bo, G. Yu, clusterProfiler 4.0: A universal enrichment tool for interpreting omics data. Innovation (Camb) 2, 100141 (2021).

